# B cell stimulation changes the structure and higher-order organization of the inactive X chromosome

**DOI:** 10.1101/2025.01.30.635789

**Authors:** Isabel Sierra, Natalie E. Toothacre, Robin H. van der Weide, Claudia D. Lovell, Son C. Nguyen, R. Jordan Barnett, Ashley L. Cook, Han-Seul Ryu, Sarah Pyfrom, Harrison Wang, Daniel Beiting, Jennifer E. Philips-Cremins, Eric F. Joyce, Montserrat C. Anguera

**Author notes:** Present Address: Hubrecht Institute, Royal Netherlands Academy of Arts and Sciences (KNAW) & University Medical Center Utrecht; Utrecht; the Netherlands. Present Address: Oncode Institute; the Netherlands (Isabel Sierra, Robin H. van der Weide). These authors contributed equally (Isabel Sierra, Natalie E. Toothacre). Twitter: @montserrat (Montserrat C. Anguera).

## Abstract

X Chromosome Inactivation (XCI) equalizes X-linked gene expression between sexes. B cells exhibit dynamic XCI, with Xist RNA/heterochromatic marks absent on the inactive X (Xi) in naive B cells but returning following mitogenic stimulation. The impact of dynamic XCI on Xi structure and maintenance was previously unknown. Here, we find dosage compensation of the Xi with state-specific XCI escape genes in naive and *in vitro* activated B cells. Allele-specific OligoPaints indicate similar Xi and Xa territories in B cells that are less compact than in fibroblasts. Allele-specific Hi-C reveals a lack of TAD-like structures on the Xi of naive B cells, and stimulation-induced alterations in TAD-like boundary strength independent of gene expression. Notably, *Xist* deletion in B cells changes TAD boundaries and large-scale Xi compaction. Altogether, our results uncover B cell-specific Xi plasticity which could underlie sex-biased biological mechanisms.

## INTRODUCTION

X Chromosome Inactivation (XCI) equalizes X-linked gene expression between the sexes in female placental mammals. XCI occurs during early embryonic development through a multistep process initiated by the expression of the long non-coding RNA Xist^1–4^. The choice of X for silencing in XCI is random, and the future inactive X chromosome (Xi) can be identified by upregulation and spreading of Xist RNA transcripts. Various repressive histone tail modifications and DNA methylation are then added to the Xi, resulting in gene repression across most of the Xi^5–11^. The enrichment of Xist RNA and heterochromatic modifications across the Xi maintain a memory of transcriptional silencing with each cell division that is maintained into adulthood. While most of the Xi is transcriptionally silenced, some genes are expressed from the Xi and thus ‘escape’ XCI^12,13^. While recent studies have identified XCI escape genes across various mouse and human tissues^14,15^, escape genes responsible for sex biased function, particularly in immune cells^16,17^ that could contribute to female-biased autoimmune diseases, are not well defined.

While the active X (Xa) retains typical features of mammalian chromosomes, including A (open/active chromatin) and B (closed/repressed chromatin) compartments, the Xi lacks compartments and TADs except at regions of gene escape as shown by Hi-C generated allele-specific spatial proximity profiles from mammalian fibroblasts and neural progenitor cells (NPCs)^18,19^. However, the presence of TADs on the Xi remains unresolved, as higher resolution Hi-C sequencing revealed global faint, low-resolution TADs^20,21^. The large-scale organization of the Xi allele is also distinct from Xa, with a unique bipartite organization involving two ‘mega-domains’ separated by the microsatellite repeat *Dxz4* region^18,19,22^. Previously the Xi was suggested to be more compacted than the Xa^23,24^, however microscopy studies have shown that the difference is fairly minimal (1.2x difference) and likely the highly spherical shape of the Xi compared to the Xa contributed to this paradigm^25^. The unique three-dimensional structure of the Xi is dependent on Xist RNA, as *Xist* deletion in fibroblasts and NPCs results in loss of the mega-domain partition and appearance of well-defined TADs across the Xi^18,26^. Thus, Xist RNA is an important contributor for maintenance of the three-dimensional structure of the Xi in somatic cells.

Most somatic cells have persistent, cytologically visible enrichment of Xist RNA and heterochromatic modifications at the Xi^27,28^. However, female lymphocytes exhibit an unusual dynamic form of XCI maintenance. In naïve B cells, the canonical Xist RNA ‘cloud’ is absent from the Xi in both mouse and human cells^29–31^, and Xist RNA re-localizes to the Xi following *in vitro* B cell stimulation with either CpG oligodeoxynucleotides (CpG) or lipopolysaccharide, before the first cell division^30,32^. Moreover, the heterochromatic histone modifications H3K27me3 and H2AK119Ub are not cytologically enriched on the Xi in naive B cells, and these marks appear concurrently with Xist RNA following *in vitro* B cell stimulation^30,32^. Naive B cells are quiescent, with significantly reduced transcription rates and highly condensed chromatin^33^. Yet, *Xist* is constitutively expressed in naive B cells, at similar levels as *in vitro* activated B cells where Xist RNA ‘clouds’ are localized at the Xi^30,34^. However, whether transcriptional repression is preserved across the Xi in naive B cells lacking enrichment of Xist RNA and heterochromatic histone modifications is unknown. Moreover, *in vitro* B cell stimulation causes significant genome-wide chromatin decondensation and rearrangement within the nucleus all before the first cell division^35,36^, but whether the chromosome architecture of the Xi changes with B cell stimulation, as Xist RNA and heterochromatic marks become enriched at the Xi, is unknown.

Here we examine the transcriptional activity of the Xi in naive (0 hr) and *in vitro* activated (12 hr, 24 hr post-stimulation) female mouse B cells to determine how cellular activation impacts gene expression and the nuclear structure of the Xi. We find that the Xi is mostly transcriptionally silent in naive (0 hr) B cells lacking localization of Xist RNA, and identify a total of 44 X-linked genes that are specifically expressed from the Xi in B cells across all timepoints. Using allele-specific OligoPaints and Hi-C analyses, we see that the global territory and compartmentalization of the Xi is relatively unchanged with B cell stimulation, yet we observe dynamic changes in Xi structure at the TAD level. Importantly, we show that *Xist* transcription, and perhaps Xist RNA transcripts themselves, are necessary for limiting stimulation induced changes to Xi TAD organization in B cells. Together, these findings provide the first evidence that Xi compaction/small scale organization is dynamic in lymphocytes and may influence XCI maintenance, providing a potential mechanistic explanation for the sex bias observed in many autoimmune diseases where B cells are pathogenic.

## RESULTS

### The Xi is dosage compensated in the absence of Xist RNA localization

To determine if Xi dosage compensation is impacted by the lack of Xist RNA ‘clouds’ at the Xi, we compared transcription between X alleles in female B cells across stimulation timepoints. For these studies, we used a mouse model of skewed XCI, in which a female *Mus musculus* mouse harboring a heterozygous *Xist* deletion is mated to a wild-type *Mus castaneus* male, generating F1 mice in which positive females (∼50% of females per F1 litter) have the paternal X chromosome inactivated in all cells and single nucleotide polymorphisms (SNPs) can be used to distinguish each allele (**Figure 1A - left)**. Using these F1 female mice, we prepared RNA for allele-specific RNA sequencing from naïve splenic CD23+ B cells, and B cells activated *in vitro* with CpG for 12 or 24 hrs (**Figure 1A – right)**. We also performed RNA sequencing in primary skin fibroblasts derived from the same F1 mice, which show canonical Xist RNA localization at the Xi, as a control. Principle component analysis of genome-wide gene expression across all samples revealed separation based on stimulation and cell type **(Figure S1A)**. We observe that the Xi is more transcriptionally silent than the active X (Xa) in B cells (0, 12, 24 hrs), with significantly less reads (reads per kilobase of transcript per million reads mapped (RPKM)) from the Xi compared to the Xa (**Figure 1B)**. In female fibroblasts, the majority of reads are also expressed from the Xa compared to the Xi **(Figure S1B).** Xi gene expression in B cells is detected from various regions across the X in both naive and *in vitro* stimulated B cells, with slightly higher expression of genes located at the proximal end of the chromosome compared to distal, excluding the *Xist* locus (**Figure 1C)**. Fibroblasts have similar regions containing XCI escape genes as B cells (**Figure 1C, Figure S1C)**, despite the difference in Xist RNA localization dynamics^30,34^. In B cells, the Xi and Xa also show similar regions of expression **(Figure S1D)**, but with higher levels of expression from the Xa compared to the Xi.

**Figure 1.**
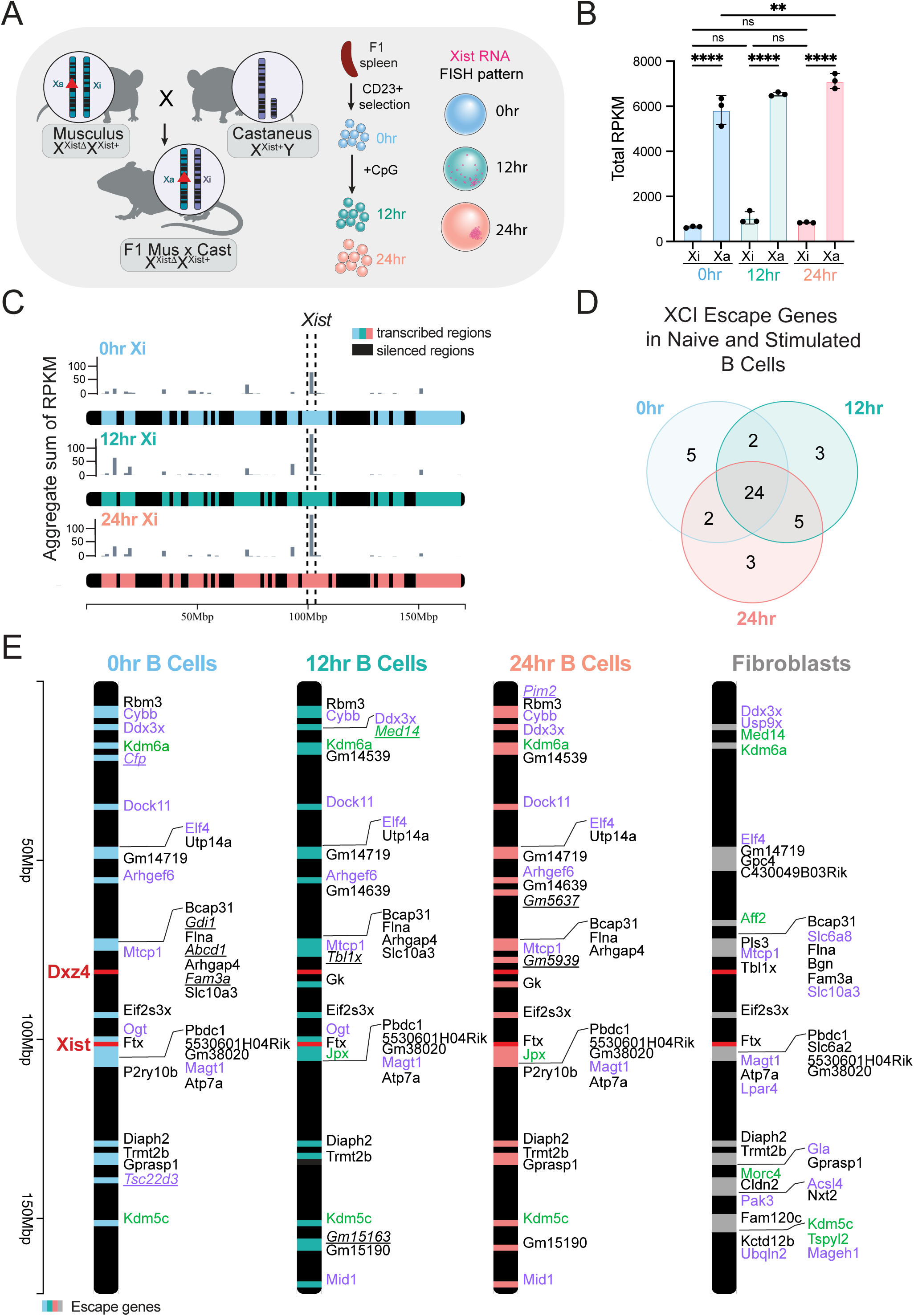
The inactive X in B cells is dosage compensated and has XCI escape genes in female B cells. (**A)** Schematic for the F1 mus x cast mouse with skewed XCI. Female *Mus musculus* (Musculus) heterozygous for *Xist* deletion (XistΔ) is mated to a wild-type male *Mus castaneus* (Castaneus) mouse. The F1 generation expresses *Xist* exclusively from the Castaneus allele, thus the wildtype Xi is paternally inherited (Castaneus) in every cell of this mouse. Splenic CD23+ B cells from female F1 mus x cast mice are stimulated *in vitro* with CpG for 12 or 24 hours, and used for the experiments in this study. Cartoon representation of Xist RNA FISH patterns at each timepoint are previously reported^30^. (**B)** Allele-specific RNAseq analyses showing total reads per kilobase per million mapped (RPKM) of X-linked reads mapping to either the Xi or Xa genomes in B cells across 0, 12, 24 hrs. Bars represent mean +/- SD. Statistical test performed using one-way ANOVA with Tukey’s correction for multiple comparisons. ** p-value < 0.005, **** p-value < 0.0001, ns = not significant. (**C)** X chromosome plots generated by chromoMap^55^ showing the genomic location of Xi-specific expression in 0hr (blue), 12 hr stimulated (green), and 24 hr stimulated (coral) B cells, for one replicate sample. Values are displayed as the sum of RPKM in 1.7Mb bins. *Xist* locus shown with dotted lines. Colored regions represent annotated and expressed regions of the X chromosome, black bars represent silenced and un-annotated regions. (**D)** Venn diagram showing the distribution of the 44 XCI escape genes in female B cells across timepoints. (**E)** X chromosome maps showing the location of each XCI escape gene in B cells and female fibroblasts. *Xist* and Dxz4 regions are indicated with red lines. Genes are listed by location; colors indicate role in immune processes (purple), chromatin/transcription (green). Underlined and italicized genes show unique B cell XCI escape genes expressed in each timepoint.

### B cells have X-linked genes expressed from the Xi

Using the allele-specific RNAseq datasets from female F1 mus x cast mice, we applied a previously published binomial distribution approach^14^ to identify XCI escape genes in female mouse B cells and fibroblasts. We generated an allele-specific ‘XCI escape gene calculator’ where XCI escape genes are identified using various criteria: (1) genes were considered expressed from the Xi if their diploid RPKM expression was greater than 1; (2) Xi expressed genes must be SNP-accessible, with a haploid expression greater than 2; and (3) a 95% confidence interval value greater than 0 in at least 2/3 replicates per timepoint. We identified about 350 X-linked genes expressed in B cells from both alleles across replicates and timepoints (diploid RPKM**, Supplementary Table 1-3)**, of which 33 genes escaped in 0 hr, and 34 genes escaped in the 12 hr and 24 hr samples **(Supplementary Table 5)**. For female fibroblasts, we detected expression of 480 X-linked genes (**Supplementary Table 4**), and 42 genes are considered XCI escape genes **(Supplemental Table 5)**. We identified 5 XCI escape genes unique to 0 hr B cells, and 3 genes specific to either the 12 hr or 24 hr stimulated B cells (**Figure 1D).** In comparison to fibroblasts, there are 23 XCI escape genes shared between the two cell types, with 21 and 19 genes unique to B cells or fibroblasts, respectively (**Figure 1E, Figure S2A)**. There is no significant difference in the overall levels of Xi expression across B cell timepoints, and there are similar levels of total Xi expression in fibroblasts and B cells **(Figure S2B)**. When comparing the ratio of Xi-specific reads for shared specific XCI escape gene expression between B cells and fibroblasts, some genes exhibit higher Xi expression in B cells compared to fibroblasts (*Trmt2b, Eif2s3x*, *Ddx3x*, *Bcap31*) **(Figure S2C)**. Gene ontology analyses revealed that B cell XCI escape genes are involved in XCI regulatory pathways, cell size, protein transport, positive regulation of translation, apoptotic signaling pathway, and protein stabilization (**Figure S2D**). There are 12 XCI escape genes with reported immunity-related functions in B cells, including *Cybb*, *Cfp*, *Ogt*, and *Mid1* (**Figure 1E, genes in purple)**. XCI escape genes also function in transcription (*Med14*), chromatin modifications (*Kdm6a*), and regulation of chromosome architecture (*Jpx)*^37^ (**Figure 1E, genes in green)**. Intriguingly, *Jpx* exclusively escapes in stimulated B cells, which could suggest a role for increased dosage of X-linked genes in genome-wide nuclear organization changes in female B cells following stimulation.

### The B cell Xi retains canonical structure but differs from fibroblasts

Because female B cells exhibit dynamic XCI maintenance, we next asked whether the organization of the Xi territory in B cells differs from the Xi of fibroblasts, and if B cell activation induced changes with compartments and chromosome compaction on the Xi. Using our F1 mus x cast female mice, we performed allele-specific genome-wide chromosome conformation capture (Hi-C) analyses in B cells. Compartment analysis of the contact heatmaps at 200kb resolution indicates that the Xa in 0 hr and 24 hr stimulated B cells exhibits the typical checkerboard pattern indicative of A/B compartments as observed for chromosome 13 (**Figure 2A - left side, Figure S3A)**. In agreement with recent work^38^, the Xi in 0 and 24 hr stimulated B cells also has A/B compartmentalization (**Figure 2A – right side**). However, when measuring the granularity of A/B compartments, we see that the Xi has larger sized compartments (3.4Mb for Xi vs 1Mb for Xa) and a smaller percentage of A/B compartment transitions compared to the Xa **(Figure S3B).** Thus, the Xi in B cells has A/B compartments, yet compartmentalization is attenuated in comparison to the Xa. Consistent with previous reports^18,22^, we observe partitioning of the Xi into two mega-domains centered at the *Dxz4* macro-satellite region in both 0 and 24 hr stimulated B cells, detected by the second eigenvector of principle component analysis (**Figure 2A – green arrow, Figure S3C)**. We also see that the *Dxz4* region in 24 hr stimulated B cells appears to have a more defined boundary at this region compared to 0 hr B cells (**Figure 2B - top),** which is quantified using local insulation scores at 3 resolutions revealing consistently lower local insulation scores (**Figure 2B – bottom, Figure S3D).** The cellular-activation induced change in insulation at *Dxz4* region suggests that B cell stimulation might alter boundary strength at some regions across the Xi. Thus, A/B compartments exhibit more fine-grained compartmentalization on the Xa than the Xi in B cells, and B cell activation does not significantly alter global compartment structure across the Xi.

**Figure 2.**
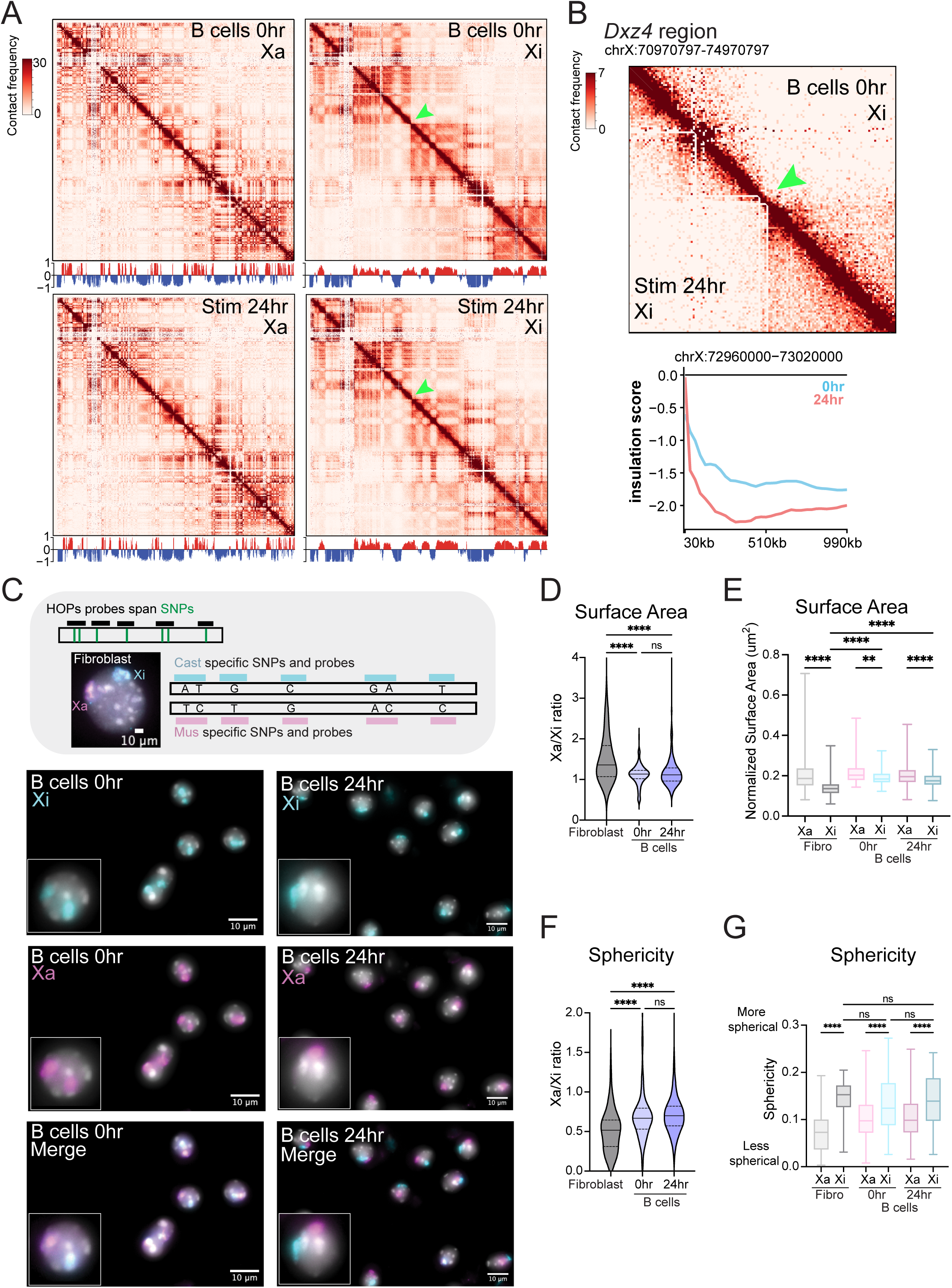
The Xi has attenuated A/B compartments in B cells and is less compact compared to primary fibroblasts. (**A)** Allele-specific Hi-C heatmaps at 200kb resolution of each X chromosome from B cells at 0 and 24 hrs post-stimulation. Green arrow denotes *Dxz4* boundary region that separates two mega-domains on Xi. Compartment tracks depicting A (red) and B (blue) compartments are below each heat map. Scale is -/+5E-2. (**B)** Hi-C heatmaps binned at 30kb resolution showing the *Dxz4* boundary region (chrX:70970797-74970797) on the Xi in B cells at 0 and 24 hrs (TOP). Quantification of the local insulation scores using sliding windows for *Dxz4* region (chrX: 72960000-73020000) on the Xi in B cells at 0 hr (blue) and 24 hr (coral) (BOTTOM). (**C)** Haplotype OligoPaints (HOPs) DNA FISH imaging for distinguishing the Xi and Xa in B cells and primary fibroblasts from female F1 mus x cast mice. Diagram of probe design for allele-specific resolution of each X chromosome. Representative DNA FISH images (single channel and merged channels) showing dual color probe labeling of the Xi (cyan) and the Xa (pink) in B cells at 0 and 24 hr timepoints. (**D)** Allele-specific surface area measurements of each X chromosome territory, calculated as the Xa/Xi ratio. **** p-value <0.0001. (**E)** Normalized allele-specific surface area measurements of each X chromosome territory, normalized by total nuclear size. ** p-value = 0.0089, **** p-value <0.0001. (**F)** Allele-specific sphericity measurements of each X chromosome territory, calculated as the Xa/Xi ratio. Violin plots show median with quartiles;**** p-value <0.0001. (**G)** Raw allele-specific measurements of sphericity for each chromosome territory. Boxplots show median with quartiles and min to max whiskers; ****p-value <0.0001. Statistical significance for surface area and sphericity measurements determined using Kruskal-Wallis tests. Hi-C experiments had n = 2 replicate female mice/timepoint. Imaging experiments had n = 3 replicate female mice for each cell type, and mice were matched for both timepoints (B cells).

As the Xi is more compact and spherical than the Xa in human diploid fibroblasts^25^, we next assessed compaction of the Xi and Xa territories in B cells using an orthogonal OligoPaints imaging method and the 3D image analysis software Tools for Analysis of Nuclear Genome Organization (TANGO) to assess allele-specific chromosome territory surface area and sphericity^39–41^ in individual nuclei. We designed a library of Homologue-specific OligoPaints for X chromosomes (HOPs-X) for DNA FISH analyses^42^, in which probes contained X-linked SNPs present in either *Mus musculus* (C57BL/6, mus) or *Mus castaneus* (cast) sequences (**Figure 2C – grey box).** Our HOPs-X probe library specifically labels each X chromosome in primary splenic CD23+ B cells (0 and 24 hr post-stimulation) (**Figure 2C)** and control primary fibroblasts isolated from F1 mus x cast mice. We observed some off-target probe binding in 0 hr B cells, thus we designed a post-TANGO image analysis processing pipeline that utilizes the integrated density of each fluorophore-labeled X chromosome to distinguish between the Xa and Xi (**Figure S4A-B, see Methods**). Quantification of the ratio of surface areas between the Xa and Xi chromosome territories in B cells revealed an Xa/Xi ratio close to 1 for both 0 hr (1.13 median) and 24 hr (1.10 median) B cells, indicating that the surface areas of the Xa and Xi are not significantly changed by activation (**Figure 2D**). The normalized surface area of the Xi is significantly smaller than the Xa in both 0 and 24 hr B cells and is not affected by B cell stimulation (**Figure 2E**). As expected, the surface area ratio (Xa/Xi) for fibroblasts is greater than 1 (1.34 median) (**Figure 2D**), yet the normalized surface area of the Xi in fibroblasts (0.13 median) is significantly lower than both the Xa and Xi in B cells (**Figure 2E, Figure S4C-D**).

As increased sphericity correlates with chromosome compaction^43^, we also measured the sphericity of the Xa and Xi chromosome territories in B cells using HOPs-X probes. We found that the sphericity Xa/Xi ratios for 0 and 24 hr B cells are similar (median naive = 0.66, median stimulated = 0.69), and significantly higher than the ratio in fibroblasts (median = 0.52) (**Figure 2F, Figure S4E-F**). Sphericity measurements revealed that the Xi territory is consistently and significantly more spherical than the Xa territory in both B cells and fibroblasts (**Figure 2G**). B cell activation causes a slight but insignificant increase in Xi sphericity (median Xi naive = 0.12, median Xi stimulated = 0.14) but no change to the Xa (median Xa naive = 0.097, median Xa stimulated = 0.098) (**Figure 2G**, **Figure S4E-F**). Together, HOPs-X measurements of the Xi and Xa territories demonstrate that the Xi is more compact and spherical than the Xa in B cells and that the organization of the Xi territory in B cells is distinct from the Xi in fibroblasts.

### The Xi gains local structures in stimulated B cells

In NPCs and fibroblasts, the Xi lacks TADs except at regions containing XCI escape genes^18,44^. Because we identified that B cell activation increased insulation of the Xi mega-domains at the *Dxz4* boundary (**Figure 2B)**, we asked whether the Xi has additional insulated structures in B cells and whether these change with B cell activation. Using our allele-specific Hi-C heatmaps binned at 30kb resolution for 0, 12 and 24 hrs stimulation, we see that there are minimal folding patterns indicative of TADs across the Xi at 0 hrs, including regions of XCI escape genes (**Figures 3A -C**). At the *Xist* region, which has the highest transcriptional level across the Xi, there are minimal TAD-like structures at 0 hrs (*Xist*, **Figure 3C**). As expected, TADs are observed across the Xa, indicating that the lack of resolvable TADs on the Xi is not a general feature of chromosomes in unstimulated B cells (**Figure S5A - C**). In 12 hr stimulated B cells, the Xi has visually more local structure, even at transcriptionally silent regions, which become stronger at 24hrs (**Figure 3A-B, green arrowheads**). The increase in local structure is reflected by the change in insulation score at 24 hrs compared to 0 hrs at these regions (**Figure 3A-B)**. To interrogate TAD-like changes chromosome-wide, we used Chromosight to identify insulated boundary regions as we were unable to call TADs on the Xi using conventional methods. Using the ‘border-detection’ function of Chromosight **(see Methods)**, we identified 206 boundaries on the Xi at 24 hrs which collectively show a slight decrease in insulation score compared to 0 hrs **(Figure S5D)**, suggesting stimulation-induced strengthening of boundary regions occurs across the Xi. As previous work indicates a link between XCI escape gene expression and TAD-like structures on the Xi^18^, we investigated this relationship on the Xi in B cells. We computed the insulation score for promoters of the 44 XCI escape genes compared to repressed genes on the Xi (n=248) (**Figure 3D)**. Promoters of XCI escape genes are associated with both insulated and non-insulated regions on the Xi when compared to silent genes, and this association is not seen on the Xa. Moreover, when taking a random sampling of the control genes to equalize gene number to escapees (n=44), we find that XCI escape gene promoters reside in closer proximity to boundaries than silenced gene promoters on the Xi **(Figure S5E)**. These findings suggest that a proportion of XCI escape genes reside at insulated boundaries in B cells. Taken together, there are TAD-like structures which appear across the Xi during stimulation independent of gene expression, and XCI escape genes reside close to boundary regions.

**Figure 3.**
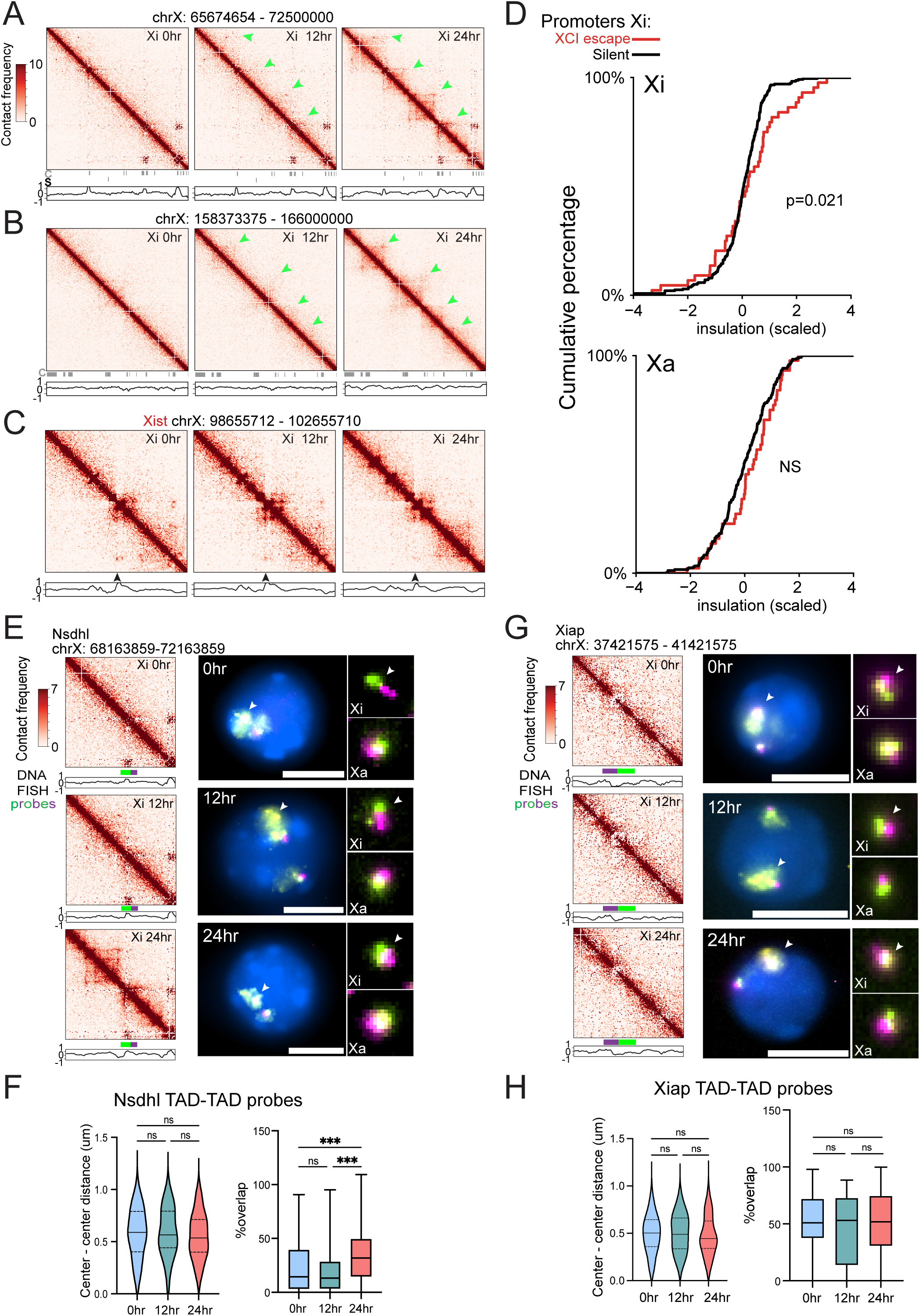
B cell stimulation increases TAD structures across the Xi. (**A)** Xi-specific Hi-C heatmaps at 30kb resolution of a region (chrX: 65674654 – 72500000) in B cells across 0, 12, 24 hrs timepoints. Green arrows denote increased TAD interactions as stimulation progresses. Insulation scores (green line) are shown below each heat map, along with locations of repressed genes on the Xi (C) and XCI escape genes (S). (**B)** Xi-specific Hi-C heatmaps at 30kb resolution of a region (chrX: 158373375 – 166000000) in B cells across 0, 12, 24 hrs timepoints. Insulation scores (green line) are shown below each heat map, along with locations of repressed genes on the Xi (C). (**C)** Xi-specific Hi-C heatmaps at 30kb resolution of a 4Mb region (chrX: 98655712 – 102655712) encompassing the *Xist* gene (black arrowhead) in B cells across 0, 12, 24 hrs timepoints. Insulation score is shown below each heatmap. (**D)** Insulation score (median score over timepoints) distribution for promoters of all B cell escape genes (red, n=44) relative to genes subject to XCI silencing (black, n=248) for the Xa and Xi. P-value was determined using a one-sided Kolmogorov-Smirnov test and is indicated on graphs. (**E)** Detection of TADs at the *Nsdhl* region on the Xi, using Hi-C and two-color DNA FISH imaging. (*Left*) Hi-C heatmaps (30kb) of *Nsdhl* region on the Xi in B cells across 0, 12, 24 hrs timepoints, TAD probe location (green, purple) spanning the boundary and insulation score shown below each heatmap. (*Right*) DNA FISH imaging of the TAD at the *Nsdhl* regions on the Xa and Xi (white arrowhead). Representative images of single nuclei at each timepoint; zoomed panels show probe-labeled TADs for each X chromosome. (**F)** Quantification of bi-color probe overlap for *Nsdhl* TAD region on the Xi. Measurements of center to center distances between each probe color and boxplot of percent overlap for the two colors. Percent overlap is normalized to the volume of the green probe. (**G)** Imaging of the *Xiap* region, lacking TADs and spanning no boundary, on the Xi using Hi-C and two-color DNA FISH. (*Left*) Hi-C heatmaps (30kb) of *Xiap* region on the Xi in B cells across 0, 12, 24 hrs timepoints, two-color probe location (green, purple), and insulation score shown below each heatmap. (*Right*) DNA FISH imaging of the TAD at the *Xiap* region on the Xa and Xi (white arrowhead). Representative images of single nuclei at each timepoint; zoomed panels show probe-labeled regions for each X chromosome. (**H)** Quantification of bi-color probe overlap for the *Xiap* region. Measurements of center to center distances between each probe color and boxplot of percent overlap for the two colors. Percent overlap is normalized to the volume of the green probe. For DNA FISH imaging experiments, n=3 replicate female F1 mus x cast mice for 0 and 24 hrs timepoints; n=2 female mice for 12 hr timepoint. Scale bars are 5um. Statistics were performed using a Kruskal-Wallis test, *** p-value < 0.0005.

To support our Hi-C results, we used a two-color OligoPaints DNA FISH assay to quantify TAD-level changes across the Xi during B cell activation. This assay measures inter-TAD interactions of two adjacent regions through quantification of spatial distance and percent overlap, which serves as an indirect measurement of chromatin remodeling^45^. We designed oligos at specific Xi regions that contained (*Nsdhl*) or lacked (*Xiap*) an insulated boundary in stimulated B cells. For each nucleus, we calculated both the center-to-center distances and percent overlap for each pair at 0, 12 and 24 hrs post-stimulation. We paired our TAD probes and HOPs-X allele-specific probes (specific to either Xa or Xi) to identify TADs on the Xi in B cells. Hi-C heatmaps for the *Nsdhl* region (ChrX:69166360-71161360) indicate that 0 hr B cells lack TAD-like structures on the Xi (**Figure 3E – top left**). There are slight structural changes for this region on Hi-C heatmaps at 12 hrs post-stimulation and increased boundary insulation at 24 hrs post-stimulation (**Figure 3E - left).** While there is a slight decrease in center-to-center distances for the *Nsdhl* TAD probes with B cell stimulation, there is a significant increase in the signal overlap at 24 hours post-stimulation (**Figure 3E-F)**, supporting observations of the Hi-C contact matrix at 24 hrs. At 12 hrs, the percentage of probe overlap is similar to 0 hr timepoint (**Figure 3F - right**), which is in agreement with the 0 and 12 hrs Hi-C matrices showing weak contact frequencies (**Figure 3E**). For the *Xiap* region, the Hi-C heatmaps lack TAD-like structures for 0, 12, and 24 hrs (**Figure 3G**) and there are similar levels of probe center-to-center distances and probe overlap following TAD FISH quantification (**Figure 3H**). Interestingly, the same regions on the Xa follow similar trends **(Figure S5F-G)**, suggesting similar regulatory mechanisms may occur on both alleles. Taken together, B cell stimulation induces remodeling across the Xi at 24 hrs post-stimulation.

### Xist deletion alters Xi structure in female B cells

Xist RNA transcripts are localized to the Xi in B cells between 12 and 24 hrs post-stimulation^30^, thus we asked whether Xist RNA affects TAD interactions on the Xi. We deleted *Xist* in B cells by mating Xist 2lox mice^46^ to Mb1-Cre recombinase animals, where Cre Recombinase is expressed starting at the pro-B cell stage and deletion is maintained in CD23+ B cells^47^. *Xist* is expressed in all B cell progenitors despite the lack of Xist RNA localization at the Xi in B cell progenitors^30^. Splenic CD23+ B cells from female wildtype and homozygous *Xist^cKO/cKO^*(*Xist^cKO^*) mice were isolated and stimulated *in vitro* for 24 hours (**Figure 4A)**. Stimulated B cells from *Xist^cKO^* mice lack detectable Xist RNA ‘clouds’ as analyzed by Xist RNA FISH (**Figure 4B)**. We performed RNA sequencing of splenic B cells (both 0 and 24 hrs) from female *Xist^cKO^* and wildtype female littermates and observed that 0 hr samples clustered together and apart from 24hr samples (**Figure S6A**). The *Xist^cKO^* 0 hr B cells have significant overexpression of two X-linked genes (Cfp, Ftx) yet 24 hr B cells do not have any differentially expressed X-linked genes (**Figure S6B**). *Xist* deletion also impacts autosomal gene expression in B cells, where 0 hr have more differentially expressed genes (∼95 genes) compared to 24 hr (∼18 genes) (**Figure S6C**).

**Figure 4.**
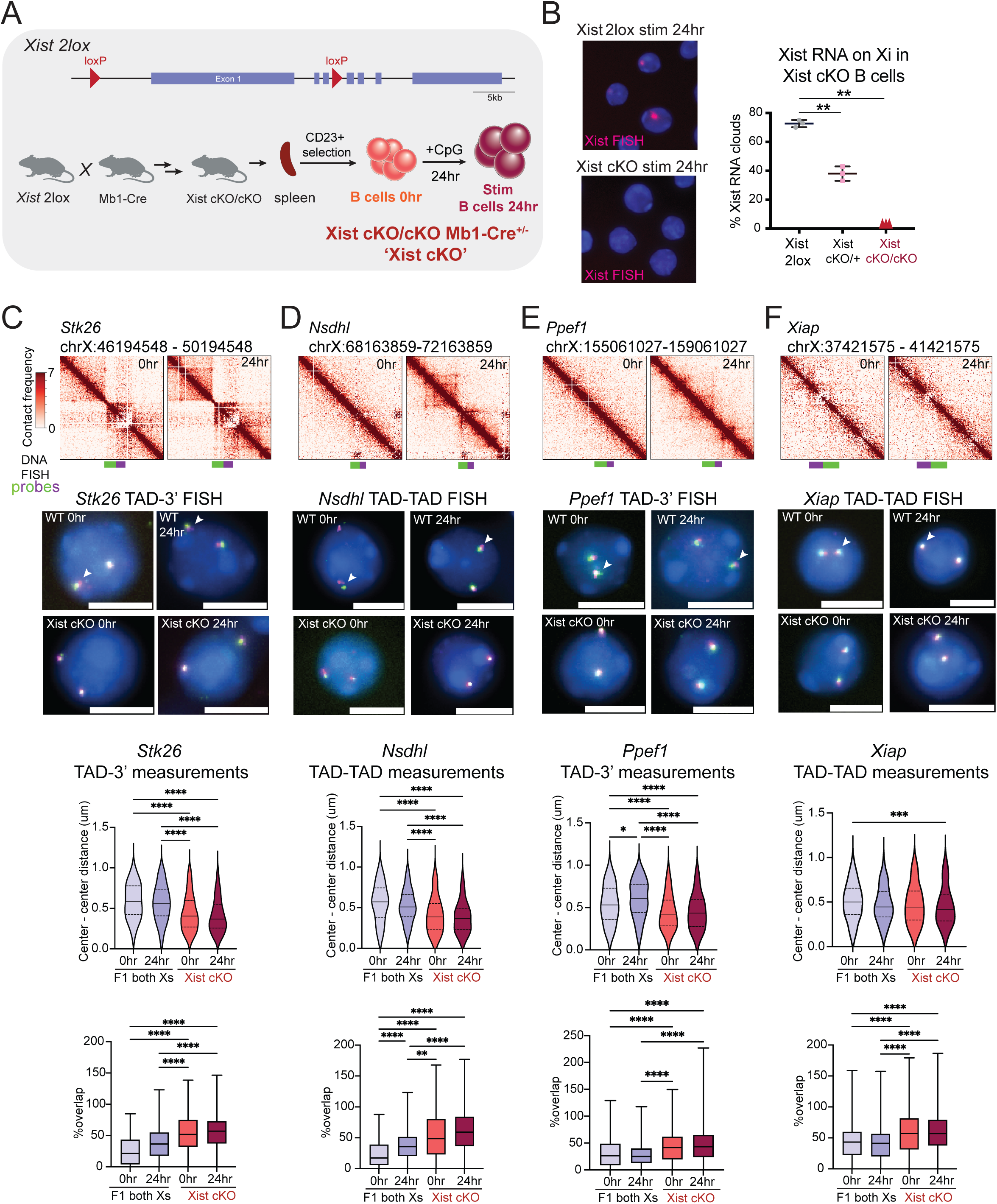
*Xist* deletion in B cells increases inter-TAD interactions across the Xi. **(A)** Schematic of the *Xist* locus and loxP sites in *Xist2lox* mice for *Xist* deletion in mature CD23+ B cells using Mb1-Cre recombinase to generate *Xist cKO/cKO (‘Xist cKO’)* mice after multiple mating steps. See Methods for further details. (**B)** Representative Xist RNA FISH images of *in vitro* activated B cells (24hrs) from *Xist2lox* and *Xist cKO* mice showing loss of Xist RNA signal; quantification of nuclei with robust Xist RNA clouds for *Xist2lox*, *Xist cKO/+* and *Xist cKO* mice (n = 3 for each genotype). Significance was determined using unpaired t-test, ** p-value <0.001. **(C-F)** Detection of TADs across the X-chromosomes, using allele-specific Hi-C and two-color DNA FISH imaging, for B cells at 0 and 24 hrs timepoints. DNA FISH imaging of TADs using 2-color probes (green, magenta) measured either TAD-TAD boundaries or TAD-3’ boundary region, centered (+/-2 Mb) on the (**C)** *Stk26 region*; (**D)** *Nsdhl* region; (**E)** *Ppef1* region; (**F)** *Xiap* region (lacking TADs or Xi boundary, control). Images show representative nuclei for Xist 2lox (F1 mus x cast) and Xist cKO B cells at each timepoint, and white arrowheads indicate the Xi in Xist 2lox samples (Xi cannot be distinguished from Xa in Xist cKO samples). Scale bars for TAD imaging are 5um. Quantification of 2-color probe overlap (center to center distances and probe color overlap) for each region is shown below. Percentage of probe color overlap is normalized to the volume of the green probe within in each pair. Statistical significance determined using a Kruskal-Wallis test; *p-value < 0.05, **p-value < 0.005, ***p-value = 0.0006, ****p-value < 0.0001. n = 3 replicate female mice for each genotype; n = 2 replicate female F1 mice for *Stk26* region at 24hrs.

Using female *Xist^cKO^* mice, we performed 2-color TAD FISH assays designed to span Xi boundaries to quantify inter-TAD interactions, examining one region with TAD-like structures in both 0 and 24 hrs B cells (*Stk26) (***Figure 4C**), two regions where TAD structures emerged at 24 hrs in Hi-C heatmaps (*Nsdhl, Ppef1*) (**Figure 4D, 4E**), and one region that lacked TADs in both 0 and 24 hrs timepoints (*Xiap*) and does not span an Xi boundary (**Figure 4F**). Of note the *Stk26* and *Ppef1* probes measure a TAD and an intervening region 3’ to the green TAD probe. We measured the center-to-center distances and amount of probe overlap at each region in *Xist^cKO^* B cells at 0 and 24 hrs, and compared measurements to the combined Xi and Xa results from our F1 B cells to assess TAD remodeling activity. For *Stk26, Nsdhl*, and *Ppef1* regions, we found that *Xist^cKO^* naive B cells have significantly shorter center-to-center distances and greater probe overlap compared to wildtype B cells at 0 and 24 hrs (**Figure 4B-D**), reflecting increased TAD interactions. B cell activation itself did not significantly change center-to-center probe distances or overlap for *Xist^cKO^* B cells (**Figure 4B-D**). At the *Xiap* locus that lacks detectible TADs at both timepoints in wildtype B cells, *Xist* deletion slightly reduced the center-to-center distances and increased the percent probe overlap relative to wildtype B cells (**Figure 4E**). Next, we examined the absolute difference of both the distance and percent overlap measurements between the X chromosome alleles when *Xist* is deleted **(Figure S7A-D)**. We see that absolute difference of distance does not significantly change after loss of *Xist*, and there are minimal changes with percent overlap differences between X alleles, suggesting that *Xist* deletion does not alter the magnitude of variation between the Xa and Xi. As differences in large-scale compaction of the Xa and Xi are more evident than differences at the TAD scale^48^, we wondered if *Xist* deletion altered sphericity of the Xi. Using non-allele specific Oligopaints, we measured the sphericity of each X allele in Xist cKO 0hr and 24hr B cells and calculated the absolute difference of X alleles **(Figure S7E)**. We see that in wildtype B cells (F1 cells), sphericity of X alleles becomes more similar after stimulation in agreement with the change in Xa/Xi sphericity ratio after stimulation (**Figure 2F)**. However, after deletion of *Xist* we no longer observe the increase in similarity between the X alleles after stimulation **(Figure S7E)**. We also asked whether occupancy of the cohesin loader protein NIPBL might be affected by *Xist* deletion. We performed CUT&RUN for NIPBL using male and female Xist^cKO^ and WT B cells, and did not see significant differences with X-linked occupancy, nor enrichment at Xi boundaries in Xist cKO B cells **(Figure S7F-H).** We observe that NIPBL occupancy increases genome-wide following B cell stimulation, and this enrichment is not specific to the X chromosome **(Figure S7H -top)**. Thus, *Xist* deletion in female B cells alters Xi structure as evidenced by changes to both large-scale compaction and smaller-scale inter-TAD interactions.

## DISCUSSION

Unlike most somatic cells, B cells utilize dynamic XCI maintenance mechanisms for dosage compensation on the Xi. During this process, Xist RNA and heterochromatic modifications, absent in naive B cells (0 hrs), localize to the Xi following mitogenic stimulation prior to the first cell division (24 hrs). How the cytological absence of Xist RNA and heterochromatic histone tail modifications in naive B cells and their recruitment to the Xi following B cell activation impacts either gene expression or Xi chromatin configuration has remained unclear. Using various allele-specific approaches to identify gene expression from the Xi and assess compaction and the presence of TAD-like structures, we provide the first evidence that Xi territory compaction and small-scale organization across the Xi is uniquely maintained in female B cells, which could potentially influence mechanisms underlying female-biased autoimmune disease.

The Xi is largely dosage compensated in naïve B cells, yet 44 X-linked genes are expressed from the Xi. These include the immunity-related genes *Cfp, Cybb, and Elf4*, raising the intriguing possibility that increased expression of X-linked immunity genes specifically from the Xi might contribute towards loss of B cell tolerance in female-biased autoimmune diseases. Although the autoimmune-associated gene *TLR7* does escape XCI in some human B cells^34,49^, we did not detect significant expression of *Tlr7* from the Xi in either 0, 12, or 24 hrs post-stimulation. However, the sustained stimulation-specific expression of the gene *Jpx* from the Xi is particularly interesting due to its recently identified role in regulating chromatin loops via CTCF^37^. *Jpx* expression coincides with the appearance of TAD-like structures across the Xi, thus it is intriguing to consider that XCI escape of X-linked chromatin regulators contribute to genome-wide chromatin reorganization that occurs in activated B cells and during their differentiation^36,50^. We detected 3-5 timepoint-specific XCI escape genes in B cells, however because we calculate Xi escape gene expression in relation to Xa gene expression, large changes in Xa gene upregulation could influence identification of XCI escape genes.

Using allele-specific Hi-C to analyze Xi structure, we found that the Xi has courser grained and attenuated compartments in B cells across all timepoints as compared to the Xa, and has the Xi-specific two mega-domains, as previously reported^18,21,22^. The Xi in B cells and neural progenitor cells (NPCs) both have compartment-like structures^38^, and compartmentalization does not change with B cell stimulation despite stronger boundary insulation between the two mega-domains after activation. The Xi in B cells is less compact compared to fibroblasts, assessed by both surface area and sphericity, which could suggest that Xi silencing in B cells is less tightly regulated than in other cell types. Possibly, the reduced compaction of the Xi in B cells allows for increased transcriptional magnitude of individual Xi escape genes in response to immune stimuli.

The Xi is quickly remodeled in B cells, where TAD contacts on the Xi increase across the Xi at the 12 and 24 hrs timepoints, with increased boundary insulation at 24 hrs. In support, the 2-color TAD DNA FISH experiments indicate that TAD-like structures are remodeled at the *Nsdhl* region on the Xi during stimulation. Previously, we have shown that the heterochromatic modifications H3K27me3 and H2AK119-ubiquitin also appear cytologically on the Xi at 12 – 24 hrs post-stimulation^30^, which coincide with the appearance of TAD structures and interactions. Additional work profiling the accumulation of these heterochromatic modifications at regions where TAD structures appear on the Xi following stimulation is necessary.

XCI escape genes are typically enriched for TAD structures^18,51^ and some B cell XCI escape genes are enriched at Xi boundary regions. However, there are increased TAD-like interactions at repressed genes on the Xi in stimulated B cells. Thus, in stimulated B cells, TADs and TAD boundary remodeling activity on the Xi does not strictly correlate with gene expression. Because B cells at 24 hrs have not yet divided, it is possible that TAD interactions will increase after cell cycle re-entry, and there might be a positive correlation between TADs and XCI escape genes at later timepoints following B cell stimulation. Additional work is necessary to determine how TAD interactions change across the Xi at timepoints beyond 24hrs, and also whether *in vivo* B cell activation changes the TAD numbers and strength across the Xi. Future studies examining how abnormal overexpression of X-linked genes and loss of Xist RNA re-localization in autoimmune disease^32,52^ impacts TAD remodeling across the Xi may provide additional insight into the relationship between transcription and chromatin organization on the Xi.

*Xist* deletion in embryonic stem cells, NPCs, and fibroblasts reconfigures the Xi chromosome^18,26^. Using our 2-color TAD imaging assay, we found that *Xist* deletion in B cells increases TAD interactions across the X, with greater overlap and shorter distances between boundary-spanning probes at all four X-loci examined. B cell stimulation did not further alter TAD interactions in *Xist^cKO^* B cells, suggesting that the loss of *Xist* transcription, instead of Xist RNA transcripts tethered at the Xi, disrupts the organization across the Xi where cellular activation does not have an additional impact on TAD interactions. Because Xist RNA is expressed but not localized to the Xi during B cell development, it is possible that *Xist* might regulate Xi organization through mechanisms independent of direct RNA tethering to regions of the Xi. Thus, we envision two possible models for *Xist*-mediated regulation of Xi compaction and TAD formation in B cells: 1. *Xist,* along with heterochromatic histone tail modifications across the Xi, are necessary for maintaining Xi compaction and limiting TAD interactions across the Xi, independent of cellular activation; 2. Upon B cell activation, Xist RNA transcripts likely recruit RNA binding proteins, heterochromatin factors, and/or architecture regulators to maintain the unique 3D compartment and specialized organization of the Xi^53,54^. Future work is needed to determine the epigenomic consequences of *Xist* deletion across the Xi in B cells. In conclusion, we propose that the Xi undergoes rapid structural changes with B cell stimulation, and that Xist RNA contributes to inter-TAD interactions across the Xi.

## LIMITATIONS OF THE STUDY

Here we used an F1 mouse model with skewed X-Chromosome Inactivation to determine 3D chromosome architecture of the inactive X in female B cells, using Hi-C and DNA FISH imaging. This study examined *in vitro* activated B cells at two timepoints after stimulation (12, 24 hrs), which is before the first cell division. Our study examined Xi compaction following cellular activation, yet it is unknown whether multiple cell divisions or B cell differentiation will impact global organization across the Xi in B cells. Our allele-specific Hi-C analyses did not detect robust TAD structures across the Xi in naive (0 hrs) B cells, however it is possible that increased sequencing depth may reveal TAD structures in unstimulated B cells. In addition, our Hi-C and DNA FISH imaging experiments were performed using *in vitro* B cell stimulation with CpG, which is an efficient and robust mechanism of cellular activation distinct from *in vivo* stimulation through immunization or infection.

## RESOURCE AVAILABILITY

### Lead contact

Requests for further information and resources should be directed to and will be fulfilled by the lead contact, Dr. Montserrat C. Anguera.

### Materials availability

This study did not generate new unique reagents.

### Data and Code availability

- The code for escape gene analysis has been deposited at Zenodo at [DOI: 10.5281/zenodo.11205424] and is publicly available as of the date of publication.
- All sequencing data generated in this study have been deposited to the NCBI GEO database with the following accession numbers: GSE215848 [Hi-C] and GSE208393 [RNAseq, CUT&RUN, Xist cKO RNAseq], and are publically available at time of publication. Source data are provided with this paper.

## ACKNOWLEDGEMENTS

We would like to thank the PennVet CHMI for their help with sequencing; L. King for help with manuscript editing; and all members from Anguera, Joyce, and Cremins labs for helpful discussions. This research was supported by an NIH R01 AI134834, Lupus Research Alliance Target in Lupus grant (to M.C.A.), NIH 1F31GM136073-01 (to I.S.), NIGMS R35GM128903 (E.F.J.), 4D Nucleome Common Fund grants U01DA052715 U01DK127405, and R01 MH120269, 1U01DA052715, 1R01-NS114226, 1DP1MH129957 (to J.E.P-C.).

## AUTHOR CONTRIBUTIONS

Conceptualization, I.S., N.E.T., M.C.A., J.E.P-C., E.F.J.; Methodology, I.S., N.E.T., S.C.N., M.C.A., E.F.J., J.E.P-C.; Investigation, I.S., N.E.T., C.D.L.; Software, S.C.N.; Formal Analysis, I.S., N.E.T., R.H.v.d.W., H.S.R., A.L.C., R.J.B., H.W., S.P.; Resources, S.C.N., D.B.; Manuscript Writing, I.S., N.E.T, and M.C.A; Manuscript Review & Editing, I.S., N.E.T., M.C.A., S.C.N., R.J.B., A.L.C., E.F.J., J.E.P-C.; Funding Acquisition, M.C.A.

## SUPPLEMENTAL INFORMATION

Document S1: Figures S1-S7

Supplemental Table S1: Allele-specific expression of X-linked genes in 0 hr B cells

Supplemental Table S2: Allele-specific expression of X-linked genes in 12 hr stimulated B cells

Supplemental Table S3: Allele-specific expression of X-linked genes in 24 hr stimulated B cells

Supplemental Table S4: Allele-specific expression of X-linked genes in female fibroblasts Supplemental Table S5: List of escape genes identified in 0 hr, 12 hr, 24 hr, B cells and fibroblast samples

Supplemental Table S6: Alignment statistics for allele-specific Hi-C Supplemental Table S7: Genomic locations of TAD imaging probes Supplemental Table S8: Xist cKO RNAseq

## STAR METHODS

### Key Resources Table

| REAGENT OR RESOURCE | SOURCE | IDENTIFIER |
| --- | --- | --- |
| Antibodies |  |  |
| anti-CD23 (Clone B3B4) | Biolegend | 101604 |
| Anti-IgG | GenScript | Cat #A01008 |
| Anti-NIPBL antibody | Bethyl Laboratories | A301-778A |
| Purified Rat Anti-Mouse CD16/CD32 | BD Biosciences | 553142 |
| Anti-B220-BV650 (clone RA3-6B2) | Biolegend | 103241 |
| Anti-CD19-BV785 (clone 6D5) | Biolegend | 115543 |
| Anti-CD23-BV421 (B3B4) | Biolegend | 101621 |
| Anti-CD43-PE (S11) | Biolegend | 143205 |
| Bacterial and virus strains |  |  |
| Biological samples |  |  |
| Chemicals, peptides, and recombinant proteins |  |  |
| CpG | Invivogen | tlrl-1826 |
| streptavidin microbeads | Miltenyi | 130048101 |
| TRIzol reagent | ThermoFisher | 15596026 |
| concanavalin A-coated beads | Polysciences | Cat #86057-3 |
| PIC | Roche | Cat #11697498001 |
| RNase A | Roche | Cat #3335399001 |
| Proteinase K | Thermo Scientific | Cat #EO0491 |
| magnetic LS column | Miltenyi | 130042401 |
| Linear Acrylamide | VWR | Cat #K548 |
| Critical commercial assays |  |  |
| Illumina TruSeq Stranded Total RNA Stranded Ligation Ribo-Zero+ kit | Illumina | 20040529 |
| Arima Hi-C+ kit | Arima Genomics |  |
| KAPA HyperPrep kit | Roche | 07962312001 |
| NEBNext Ultra II DNA Library Prep Kit for Illumina | NEB | Cat # E7645S |
| LIVE/DEAD Fixable Aqua Dead Cell Stain Kit | Life Technologies | L34965 |
| Qiagen RNeasy Micro Kit | Qiagen | 74004 |
| SMARTer Stranded Total RNA-Seq Kit v3 - Pico Input Mammalian | Takara | 634485 |
| Deposited data |  |  |
| B cell Hi-C data | National Center for Biotechnology (NCBI) | GSE215848 |
| B cell RNA sequencing, CUT&RUN, Xist cKO RNA sequencing | National Center for Biotechnology (NCBI) | GSE208393 |
| Experimental models: Cell lines |  |  |
| Experimental models: Organisms/strains |  |  |
| Xist<sup>fl/fl</sup> female mice | Jackson Labs | strain# 000664 |
| Mus castaneus | Jackson Labs | strain# 000928 |
| B6.C(Cg)-Cd79a<sup>tm1(cre)Reth</sup>/Ehobj | Jackson Labs | strain# 020505 |
| B6N.FVB-Tmem163<sup>Tg(ACTB-cre)2Mrt</sup>/CjDswJ | Jackson Labs | strain# 019099 |
| Oligonucleotides |  |  |
| Recombinant DNA |  |  |
| Software and algorithms |  |  |
| SNPsplit |  |  |
| allele-specific Hi-C analysis using Hi-C-Pro v2.8.9 | (https://nservant.github.io/Hi-C-Pro/AS.html) |  |
| GENOVA v1.0.1 | Ref. 65 |  |
| chromosight v1.6.3 | Ref. 67 |  |
| deepTools 3.5.4 | Ref. 68 |  |
| OligoMiner design pipeline | Ref. 69 |  |
| TANGO | Ref. 70 |  |
| Other |  |  |

### Experimental Model and Study Participant Details

#### Mice

For generation of the F1 mus x cast mice with skewed XCI, *Xist^fl/fl^* female mice (‘Xist2lox’; C57BL/6j; strain# 000664, Jackson) were mated to an ACTB-Cre male (B6N.FVB-*Tmem163^Tg(ACTB-cre)2Mrt^*/CjDswJ; strain# 019099, Jackson) to generate heterozygous *Xist^fl/+^* females, which were bred with *Mus castaneus* (Cast; strain# 000928, Jackson) males to generate F1 *Xist^fl/+^* female mice. To generate the B cell specific Xist cKO knockout mice, Xist2lox mice were bred to an Mb1-Cre Recombinase mice (B6.C(Cg)-*Cd79a^tm1(cre)Reth^/*Ehobj; strain# 020505; Jackson) to generate mice heterozygous for Mb1-Cre, then intercrossed (and backcrossed for 10 generations to C57Bl/6j) to generate homozygous Xist cKO/cKO *Mb1-cre^+/-^* animals. The Mb1-Cre Xist cKO mice do not have allelically-skewed XCI as the *Xist* deletion occurs after establishment of random XCI. Therefore, heterozygous crosses will have *Xist* deleted from an inactive X in approximately 50% of B cells, while homozygous crosses have *Xist* deleted on both alleles in all B cells (100% of Xi alleles contain an *Xist* deletion). All mice were maintained at the Penn Vet animal facility, and experiments were approved by the University of Pennsylvania Institutional Animal Care and Use Committee (IACUC). Euthanasia via carbon dioxide was used for animal sacrifice prior to isolations.

#### Generation of primary fibroblast lines

Primary adult mouse fibroblasts were isolated as previously described^56^. Ears from n= 3 replicate female F1 mus (C57BL/6j) x cast mice were removed and cut into 3mm size pieces, then incubated in collagenase D-pronase solution for 90 min at 37C with agitation, then ground and filtered through a 70um cell strainer into complete medium [RPMI with 10% fetal calf serum, 50uM 2-mercaptoethanol, 100uM asparagine, 2mM glutamine, 1% penicillin-streptomycin]. Cells were washed then plated with complete medium supplemented with 10ul amphotericin B [250ug/ml stock], and cultured until they reached 70% confluency. Cells were passaged 1-2 times before collection for RNA isolation or imaging.

### Method Details

#### B cell isolations

Follicular B cell isolations were performed as previously described using a positive selection kit^30^. Briefly, spleens from mice aged 3-6 months of age were crushed to produce single cell suspensions, then incubated with Biotin tagged anti-CD23 (Clone B3B4, 101604, Biolegend) followed by streptavidin microbeads (130048101, Miltenyi). Cells were run through a magnetic LS column (130042401, Miltenyi) for positive selection of CD23+ B cells. B cells were collected immediately for experiments (0 hr timepoint) and half were cultured with 1uM CpG (tlrl-1826, Invivogen) for 12 hrs or 24 hrs timepoints.

#### RNA Isolation and Allele-specific RNAseq sequencing

B cells isolated from n = 3 replicate F1 mus (C57BL/6j) x cast female mice were collected at 0 hr, 12 hr, and 24 hr post stimulation into TRIzol reagent (15596026, ThermoFisher). Primary fibroblasts (n = 3 replicate mice) from F1 mus x cast mice were collected into TRIzol reagent. RNA isolations were performed according to the manufacturers protocol. Libraries were prepared with an Illumina TruSeq Stranded Total RNA Stranded Ligation Ribo-Zero+ kit (20040529, Illumina), pooled, and run on an Illumina NextSeq 2000 sequencer (150bp paired-end).

To quantify gene expression allele-specifically from our F1 crosses, we created an N-masked mm10 (C57BL/6j; Ensembl GRCm38) genome using SNPsplit^57^. SNPs were derived from the Castaneus genome (CAST/EiJ, accession# ERS076381, Sanger Institute)^58^. The resulting N-masked genomes were used to generate respective STAR (v2.7.1a) indexes. Reads were aligned using STAR with alignEndsType set to EndToEnd, and outSAMattributes set to NH HI NM MD to allow for compatibility with SNPsplit. Output files were then passed through SNPsplit to allow for allele-specific sorting of reads. Reads were quantified using featureCounts^59^ on both the diploid N-masked aligned and the haploid allele-specific aligned reads.

Genes that escape XCI were identified using 3 thresholds of escape as previously described^60,61^. Briefly, diploid gene expression was first calculated in RPKM (reads per kb of exon length, per million mapped reads), and genes were called as expressed if their diploid RPKM was > 1. For every X-linked gene that passed this threshold, haploid gene expression was calculated in SRPM (allele-specific SNP-containing exonic reads per 10 million uniquely mapped reads), and genes which had an Xi-SRPM > 2 were considered expressed from the Xi. Finally, a binomial model estimating the statistical confidence of escape probability was applied to the genes passing the first 2 thresholds. This model compares the proportion of Xi-specific reads to the total Xi + Xa reads and calculates a 95% confidence interval. XCI escape status requires that the 95% lower confidence limit probability is greater than 0. To calculate relative allele-specific RPKM, the diploid values were multiplied by the allelic ratio for each X allele (Xn/Xi+Xa).

Reads were graphed using Prism v9.3.1, and statistical significance was determined using one-way ANOVA with Tukey’s multiple comparison test. The R package chromoMap was used to generate chromosome maps displaying aggregate RPKMs^55^. Venn diagraph of escape genes was generated using VennDiagram package in R. GO analysis was performed using Metascape^62^. Gene annotations in **Figure 1** were performed manually, performing literature searches for each individual gene. Heatmap was generated using ggplot2 in R.

#### Hi-C sequencing

Hi-C was performed using the Arima Hi-C+ kit (Arima Genomics, San Diego, CA, USA) following their standard protocol. 2 million cells per replicate (n = 2 mice for each timepoint) were crosslinked with 2% formaldehyde for 10 mins at RT before proceeding with Hi-C. For library preparation, the KAPA HyperPrep kit (07962312001, Roche) was used with a modified protocol provided by Arima Genomics. Libraries were checked by Tapestation analysis, pooled, and run for two rounds on a NextSeq 500 with a 2×150bp read length. Two replicates were pooled to generate 400 million reads/timepoint.

#### Allele-specific Hi-C analysis

We performed allele-specific Hi-C analysis using Hi-C-Pro v2.8.9 according to the allele-specific analysis section of the Hi-C-Pro manual (https://nservant.github.io/Hi-C-Pro/AS.html). Briefly, we generated a masked mm9 genome, in which each location of strain-specific SNPs differentiating CAST and C57Bl6 strains is N-masked. First, the NCBI37/mm9 genome was downloaded from the UCSC Genome Browser (https://hgdownload.soe.ucsc.edu/downloads.html#mouse). We then downloaded a database of strain-specific SNPs for a variety of mouse strains from the Sanger Institute Mouse Genomes Project (https://www.sanger.ac.uk/data/mouse-genomes-project/)^58^. Using this information, we generated a vcf file by running the extract_snps script on Hi-C-Pro that contained SNPs that differ between CAST and C57bl6 mouse genome. We masked the reference mm9 genome at all loci of SNPs specified in the vcf file by running the bedtools maskfasta command (https://bedtools.readthedocs.io/en/latest/content/tools/maskfasta.html). Finally, we built a Bowtie index of the masked genome using the bowtie2-build indexer (http://bowtie-bio.sourceforge.net/bowtie2/manual.shtml#the-bowtie2-build-indexer).

We aligned paired-end reads from Hi-C fastq files to our masked mm9 genome using bowtie2 (global parameters: --very-sensitive -L 30 --score-min L,-0.6,-0.2 --end-to-end – reorder; local parameters: --very-sensitive -L 20 --score-min L,-0.6,-0.2 --end-to-end – reorder). We filtered unmapped reads, non-uniquely mapped reads, and PCR duplicates (Hi-C statistics shown in **Supplementary Table 6**).

We assembled raw cis contact matrices for each chromosome at two different time points for each allele at 30kb and 200kb resolution. X chromosome alleles were assigned such that Xa represented the C57Bl/6-specific allele and Xi represented the CAST-specific allele. Replicates for each condition were merged. We balanced the merged contact matrices using Knight-Ruiz balancing as previously described^63,64^. For each balanced matrix, we performed simple scalar normalization, where a simple scalar size factor for each pixel was calculated based on

#### Compartment calling

To plot A/B compartment tracks chromosome-wide, we performed eigenvector decomposition on 200 kb resolution, balanced Hi-C matrices using the compartment_score-function of GENOVA v1.0.1^65^. We verified that the sign of the compartment score was correct by comparing GC-content in A- and B-compartments.

#### Boundary strength

We calculated insulation scores chromosome-wide by applying a 360 kb window on the 30 kb resolution, balanced Hi-C data^66^ with the insulation_score-function of GENOVA v1.0.1^65^. Next, we calculated a local domainogram at the chrX: 70970797-74970797 locus using the insulation_domainogram-function of GENOVA v1.0.1. We used window-sizes of 30 to 990 kb, with a step-size of 30 kb. For the promoter-specific analyses, we computed the allelic insulation score by taking the median insulation score across timepoints at the bins overlapping the respective promoters.

We used the detection-module of chromosight v1.6.3^67^ to detect boundaries with the following parameters: max_dist: 0, min_dist: 0, max_iterations: 5, max_perc_zero: 10.0, max_perc_undetected: 75.0, min_separation: 90000, pearson: 0.15, resolution: 30000. We filtered boundaries with a qvalue >= 0.001. We aligned the insulation scores over these boundaries with the computeMatrix-module of deepTools 3.5.4^68^ in reference-point mode at a 25 kb binsize and 1 Mb flanking regions.

#### DNA FISH probe design

Allele-specific HOPs probes libraries were mined for the X chromosome (mm10 genome build) using the OligoMiner design pipeline^69^, with the −l and -L parameters set to 42 for 42-mer oligos and the -O parameter added for overlapping oligos. The resulting set of oligos were then modified to include SNPS for the Cast/EiJ or C57BL/6NJ mouse strains, downloaded from the Mouse Genomes Project (https://www.sanger.ac.uk/data/mouse-genomes-project/). Strain-specific Oligopaints were selected using a similar workflow to the HOPs pipeline^42^. Specifically, oligos were selected based on containing at least one differential SNP in the inner 32 nucleotides of each 42-mer oligo. For TAD imaging, probes were designed to either two adjacent TAD regions (as identified from our HiC domain calling results) or a TAD and a 3’ adjacent region as indicated in figure legends. TAD probe locations are provided in **Supplementary Table 7.** Oligos were purchased from CustomArray/Genscript, and probe sets were produced as described previously^40^.

#### DNA FISH with OligoPaints

Splenic CD23+ B cells from female mice (F1 experiments: n = 3 replicate female mice for 0 hr and 24 hr, n = 2 replicate female mice for 12 hr; female *Xist cKO* n= 3 mice each) were cytospun onto slides and briefly incubated in room temperature SSCT+formamide [2X SSC, 0.1% Tween-20, 50% formamide], then pre-hybridized in SSCT+formamide for 1hr at 37C. Primary probe mix [50% formamide, 1X Dextran Sulfate Mix [10% dextran sulfate, 4% PVSA, 2X SSC, 0.1% Tween-20], 10ug RNase A, 5.6mM dNTPS, 50pmol per Oligopaint probe was added to slides, sealed with rubber cement, and denatured for 30min at 80C. Slides were hybridized overnight at 37C in a humidified chamber. Next day, the slides were washed for 15min in 2X SSCT [2X SSC, 0.1% Tween-20] at 60C, 10min in 2X SSCT at room temperature, then 10min in 0.2X SSC at room temperature. Secondary probe mix [10% formamide, 1X Dextran Sulfate Mix, 10pmol per secondary probe] was added to slides, sealed with rubber cement, and incubated in a humidified chamber for at least 2hr at room temperature. Slides were washed for 5min in 2X SSCT at 60C, 5min in 2X SSCT at room temperature, then 5min in 0.2X SSC at room temperature. Slides were mounted with Vectashield and imaged on a Nikon Eclipse microscope with Z-stacks set to a 0.2um step size.

#### DNA FISH Image analyses

All images were analyzed using TANGO^70^. The following settings were used for allele-specific images and whole X imaging in *Xist* conditional knockout cells: Nuclei – pre-filter: Fast Filters 3D; Segmentation: Hysteresis Segmenter; Post-filters: Size and Edge Filter, Morphological Filters 3D (Fill Holes 2D, Binary Close). Alleles – pre-filter: None; Segmentation: Hysteresis Segmenter; Post-filters: Size and Edge filter, Erase Spots. The following settings were used for TAD imaging: Nuclei – pre-filter: Fast Filters 3D; Segmentation: Hysteresis Segmenter; Post-filters: Size and Edge Filter, Morphological Filters 3D (Fill Holes 2D, Binary Close). TADs – pre-filter: Fast Filters 3D, Gaussian Smooth; Segmentation: Hysteresis Segmenter; Post-filters: Size and Edge filter, Erase Spots.

For allele-specific X chromosome imaging, TANGO-generated raw allele-specific data files were further processed using a python script which performed two functions: 1) utilized the integrated density values to select the true allele for each genome, with the highest value for each fluorophore being assigned as “true”. The use of this method was verified by manually checking the raw images. 2) Identified cells with overlapping or touching chromosomes to exclude from this analysis as confident allelic assignment could not be determined in these instances. For surface area measurements, each allele was normalized to its respective nuclear size. To generate surface area and sphericity ratios, the active X value was divided by the inactive X value within each individual cell then graphed. For proportion plots, the allele of choice (Xn) was divided by the sum of values for both alleles for each specific measurement (Xn/Xi+Xa). Whole X imaging in *Xist* conditional knockout cells were analyzed post segmentation by taking the absolute difference of alleles in each nucleus for sphericity measurements. The same analysis was performed on the allele-specific data for comparison. Graphs were created using the ecdf function in R v4.1.1.

For all TAD imaging, measurements were filtered out if the center-to-center distance was greater than 1um (removed trans measurements between alleles), and only cells which contained two distinct alleles (two segmented objects for each probe) were used for further analysis. For allele-specific TAD imaging, whole X allele-specific probes were used in conjunction with TAD probes, but only the Xi allele was labeled with a secondary probe. The Xi-specific TADs were manually selected by proximity to the labeled Xi allele. Distances were measured using the center-to-center distance between each TAD within each allele. Overlap values were normalized to the volume of one probe from each set (indicated in figure legends). Unless specified above, all graphs were generated using Prism v9.3.1. Significance was determined by Kruskal-Wallis statistical tests performed in Prism v9.3.1.

#### CUT&RUN for NIPBL

CUT&RUN for NIPBL was performed as previously described. Briefly, 10ul/sample of concanavalin A-coated beads (Cat #86057-3, Polysciences) were equilibrated in Binding Buffer [20mM HEPES, 10mM KCl, 1mM CaCl2, 1mM MnCl2] and resuspended in Wash Buffer [20mM HEPES, 150mM NaCl, 0.5M Spermidine, 1x PIC (Cat #11697498001, Roche)]. 1 million naive or stimulated CD23+ B cells (n=3 replicated for 0 hr WT, 0 hr cKO, 24 hr WT, 24 hr cKO) were immobilized on beads by rotating for 1 hr in Wash Buffer at room temperature. Bead-bound cells were then resuspended in Antibody Buffer [20mM HEPES, 150mM NaCl, 0.5mM Spermidine, 1x PIC, 0.05% Digitonin, 2mM EDTA]. 3ug of NIPBL antibody [A301-778A, Bethyl Laboratories] or 3ug IgG [Cat #A01008, GenScript] were added. Samples were rotated at 4C for 5 hrs. Bead-bound cells were washed and resuspended in Dig-Wash Buffer [20mM HEPES, 150mM NaCl, 0.5mM Spermidine, 1x PIC, 0.05% Digitonin]. pA-MNase fusion protein [1200ng/ml; gift from Dr. K. Sarma] was added and samples were rotated at 4C for 1 hr. Bead-bound cells were washed twice followed by resuspension in Dig-Wash Buffer. pA-MNase was activated by addition of 2mM CaCl2 and cleavage was allowed to occur for 30min at 0C. The reaction was stopped by addition of one volume of 2x STOP Buffer [340mM NaCl, 20mM EDTA, 4mM EGTA, 0.02% Digitonin, 50ug/ml RNase A (Cat #3335399001, Roche), 50ug/ml Linear Acrylamide (Cat #K548, VWR), 2pg/ml Yeast Spike-In DNA]. Chromatin was released into the supernatant through shaking at 300rpm for 10min at 37C. Chromatin-containing supernatant was transferred to new tubes and incubated at 70C for 10min with 0.1% SDS and 50ug Proteinase K [Cat #EO0491, Thermo Scientific]. Isolation of DNA was performed with phenol:chloroform:isoamyl alcohol and precipitated with ammonium acetate. Libraries were prepared using 2ng of isolated DNA using the NEBNext Ultra II DNA Library Prep Kit for Illumina [Cat # E7645S, NEB] and sequenced paired-end on a 300 cycle flow cell on the NextSeq2000.

#### CUT&RUN analysis

FASTQ files were trimmed using Trimommatic v0.32 and aligned non-allele specifically to mm10 reference genome using bowtie2 and the following parameters: -q --local --very-sensitive-local --soft-clipped-unmapped-tlen --dovetail --no-mixed --no-discordant -q -- phred33 -I 10 -X 1000 and converted to bam files using samtools. Aligned reads were de-duplicated using Picard v1.141 MarkDuplicates. Blacklist filtering [ENCODE] was performed using bedtools intersect. CPM normalized bigwigs were generated using bamCoverage with chrX ignored for normalization. For spike-in normalization, FASTQs were aligned to *E.coli* genome using bowtie2 and the following parameters: --end-to-end --very-sensitive --no-mixed --no-discordant --phred33 -I 10 -X 700. For enrichment comparisons between WT and cKO samples, each individual replicate was spike-in normalized to the number of aligned *E.coli* reads per samples and scaled accordingly. Replicates were then averaged together into one merged file using deepTools bigwigAverage. Peak calling was performed using GoPeaks with default parameters on individual replicates with IgG controls (0hr IgG and 24 IgG controls were used). Final peak calls were determined by finding shared peaks across WT replicates for 0hr and 24hr samples using bedtools intersect. To generate enrichment plots for **Supplementary** Figure 7F, deepTools computeMatrix and plotHeatmap were used on the spike-in normalized, averaged bigwigs for WT peaks in either 0hr samples or 24hr samples. For **Supplementary** Figure 7G, deepTools computeMatrix and plotHeatmap were used on the spike-in normalized, averaged bigwigs for 24hr Xi boundary locations. To determine differential binding of NIPBL between WT and cKO samples **(Supplementary** Figure 7H), the DiffBind package was used and plotted in volcano plots using EnhancedVolcano from Bioconductor in R.

#### Cell sorting Xist cKO B cells

CD23+ follicular B cells were sorted from spleens of female WT and Xist cKO mice. Splenocytes were isolated via mechanical dissociation of the spleen. Cells were stained with LIVE/DEAD Fixable Aqua Dead Cell Stain Kit (Life Technologies L34965) and subsequently blocked with Purified Rat Anti-Mouse CD16/CD32 (BD Biosciences 553142). Cells were then stained in 0.5% BSA in PBS using the following fluorochrome-conjugated antibodies (clone, catalog #) purchased from BioLegend: B220-BV650 (RA3-6B2, 103241), CD19-BV785 (6D5, 115543), CD23-BV421 (B3B4, 101621), and CD43-PE (S11, 143205). Sorting was performed on a FACSAria Fusion.

#### RNA Sequencing Xist cKO B cells

RNA was isolated from sorted populations of CD23+ follicular splenic B cells using the Qiagen RNeasy Micro Kit (Qiagen 74004). Library preparation of RNA samples was performed using the SMARTer Stranded Total RNA-Seq Kit v3 - Pico Input Mammalian (Takara 634485). Pooled libraries were sequenced on a 75 cycle flow cell using an Illumina NextSeq 2000 sequencer. Reads were aligned to a C57BL/6J reference genome (GRCm38) using STAR (v2.7.1a). Read counts were calculated using STAR quantMode “GeneCounts”. RNAseq data was analyzed using RStudio (v1.2.5042). edgeR (v3.32.1) was used to filter data and data was normalized using quantile normalization. Differentially expressed genes were identified using DESeq2 (v1.30.1), and are shown in Supplemental Table 8.

### Quantification and Statistical Analyses

For TAD analyses, all graphs were generated using Prism v9.3.1. Significance was determined by Kruskal-Wallis statistical tests performed in Prism v9.3.1.

