## Supplementary figures and images for "B cell stimulation changes the structure and higher-order organization of the inactive X chromosome"

### Supplemental Figures

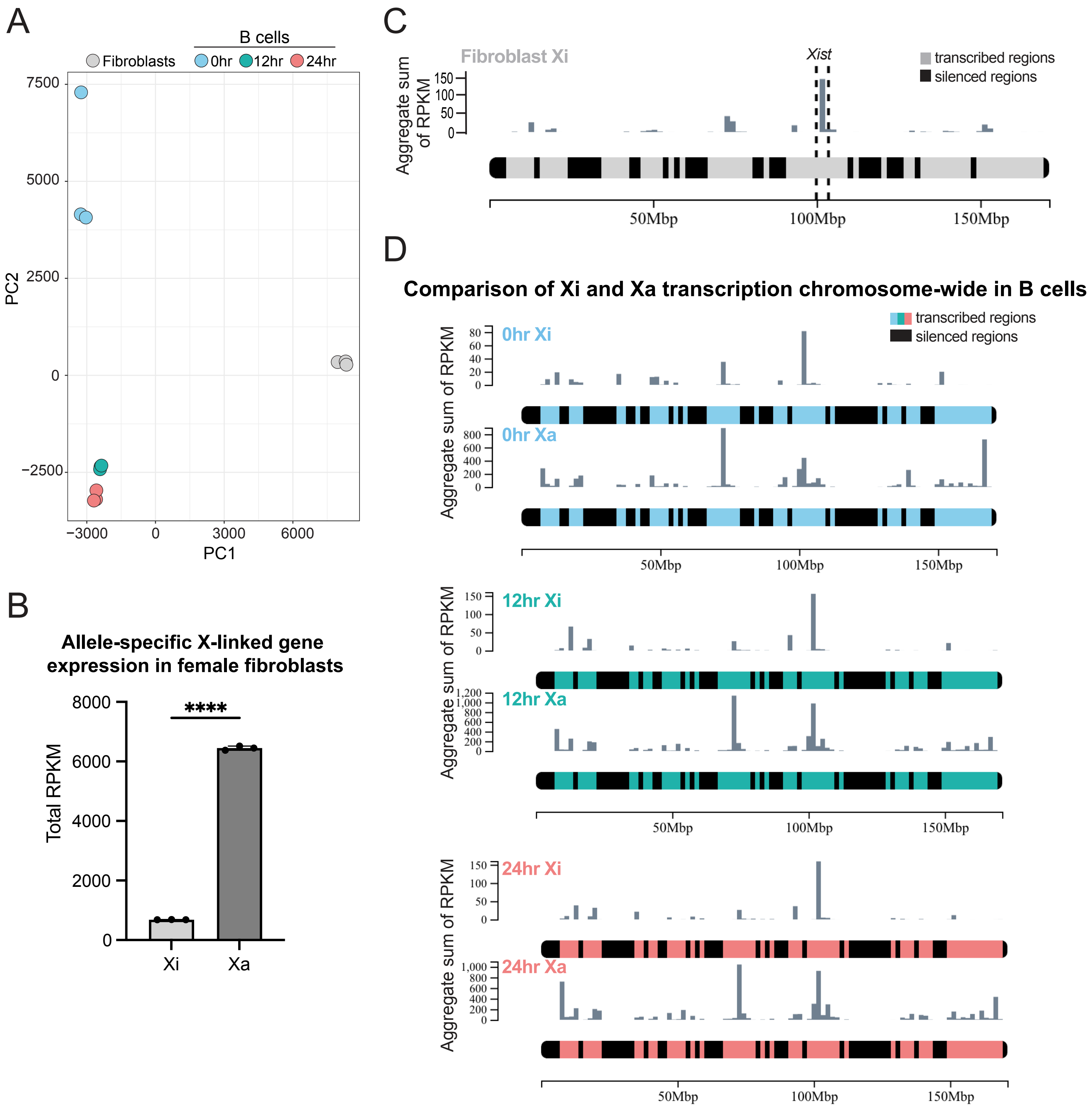

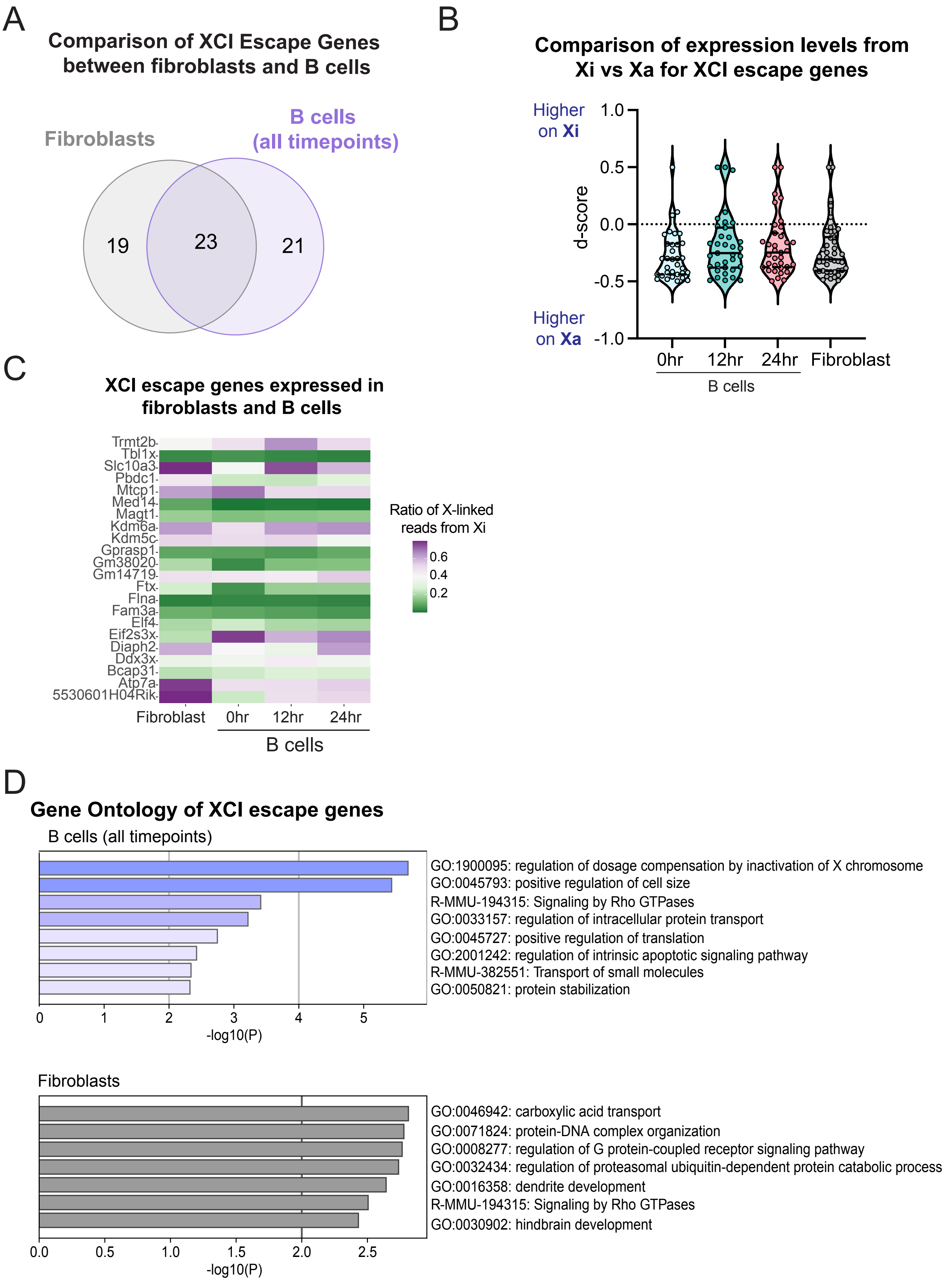

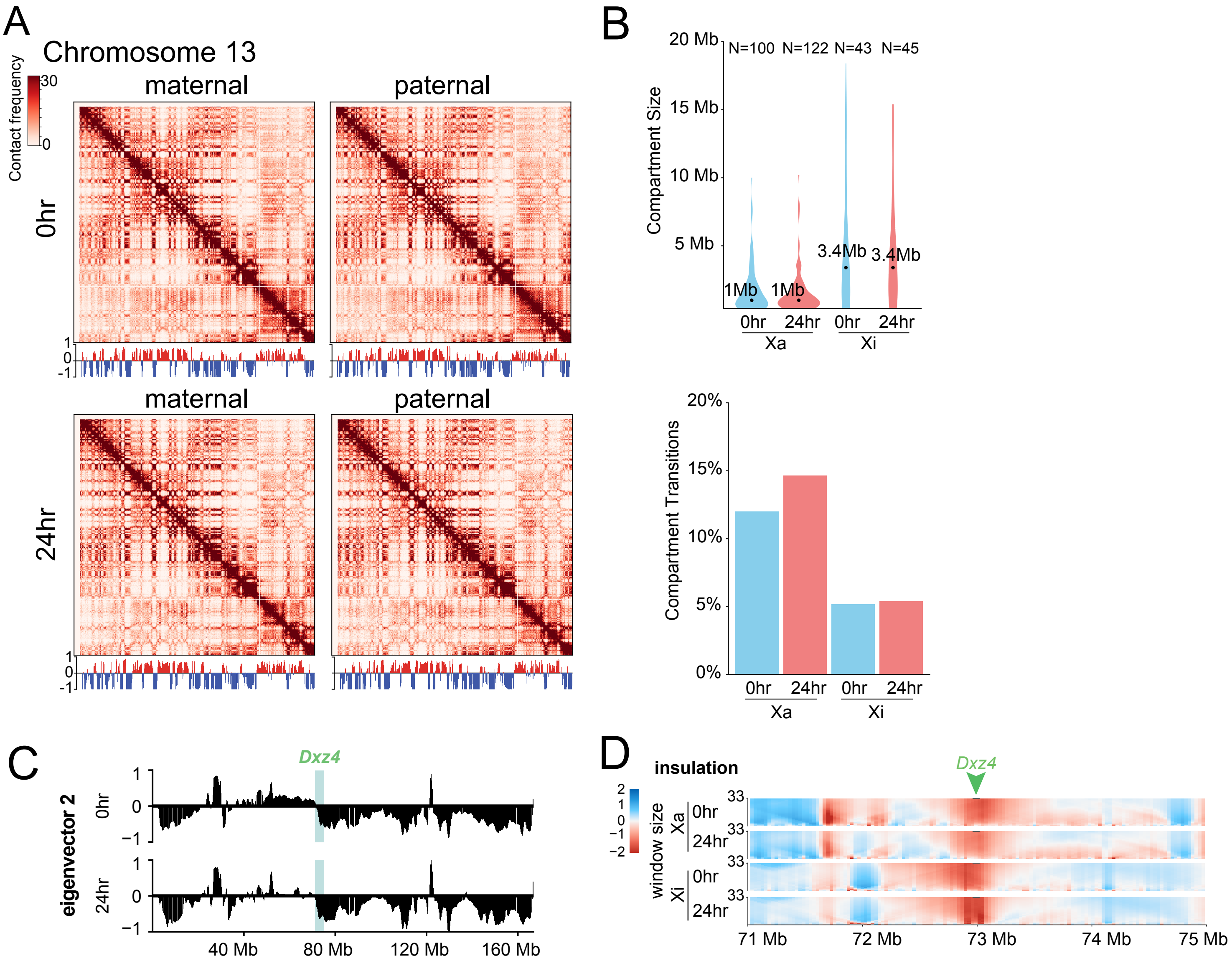

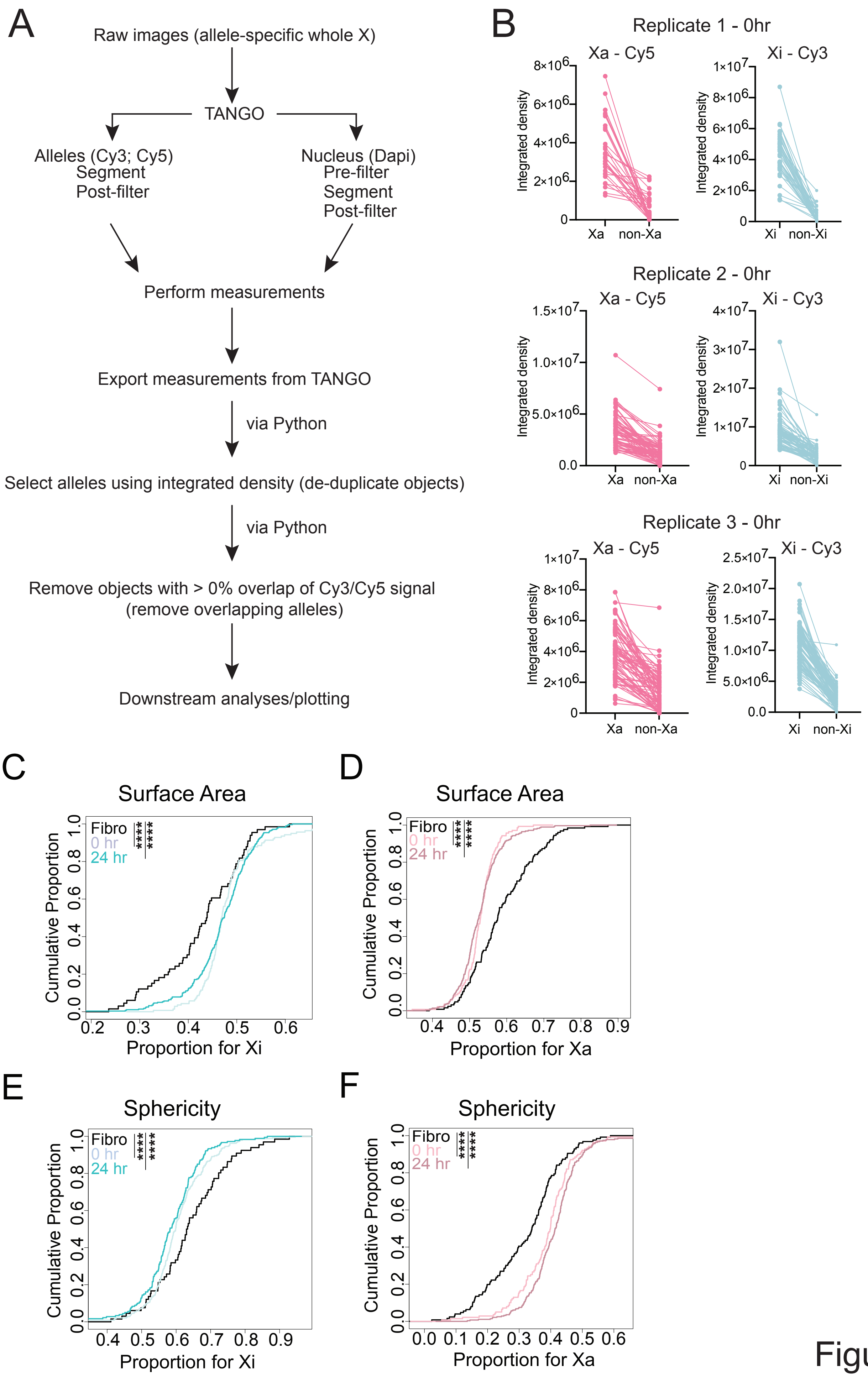

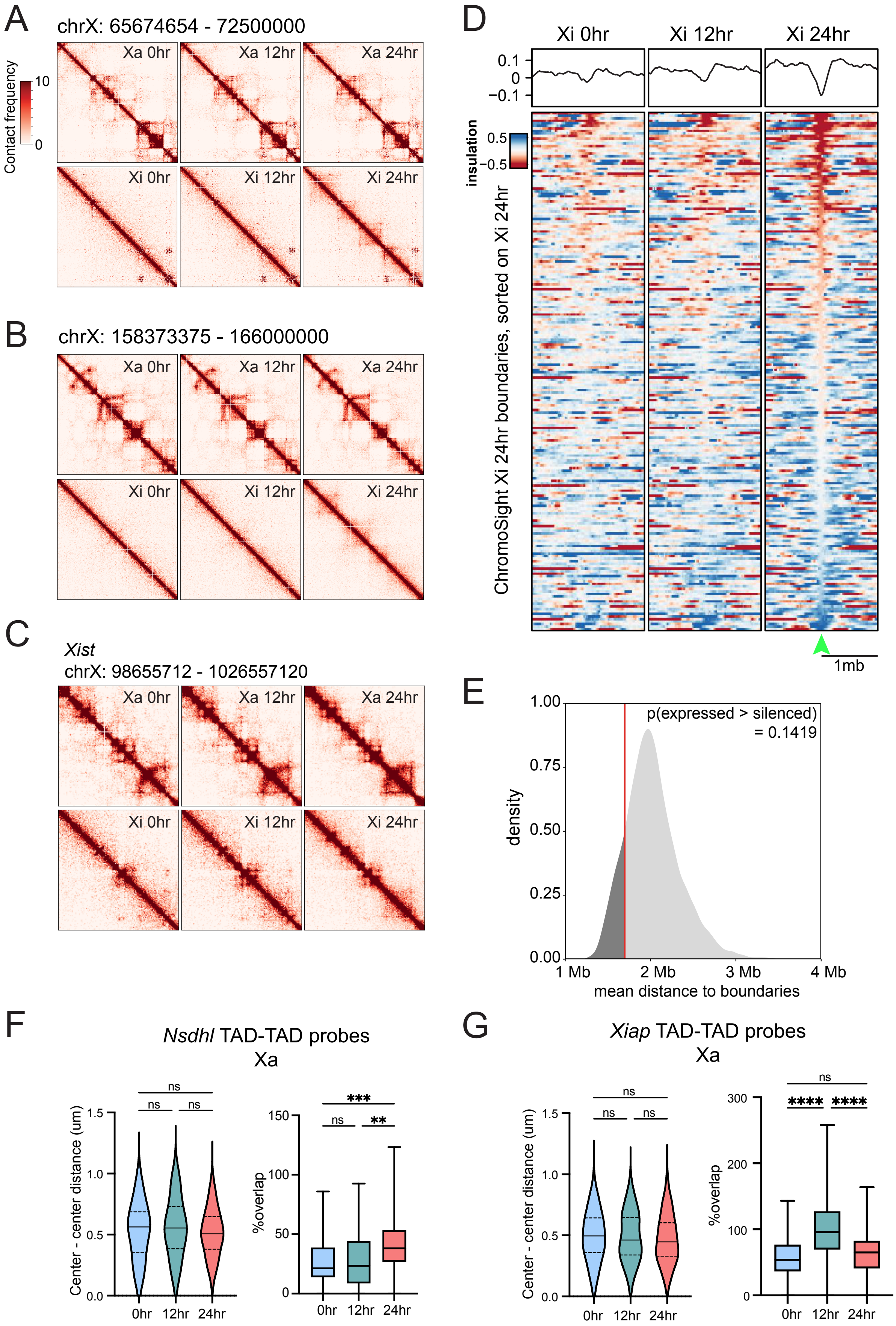

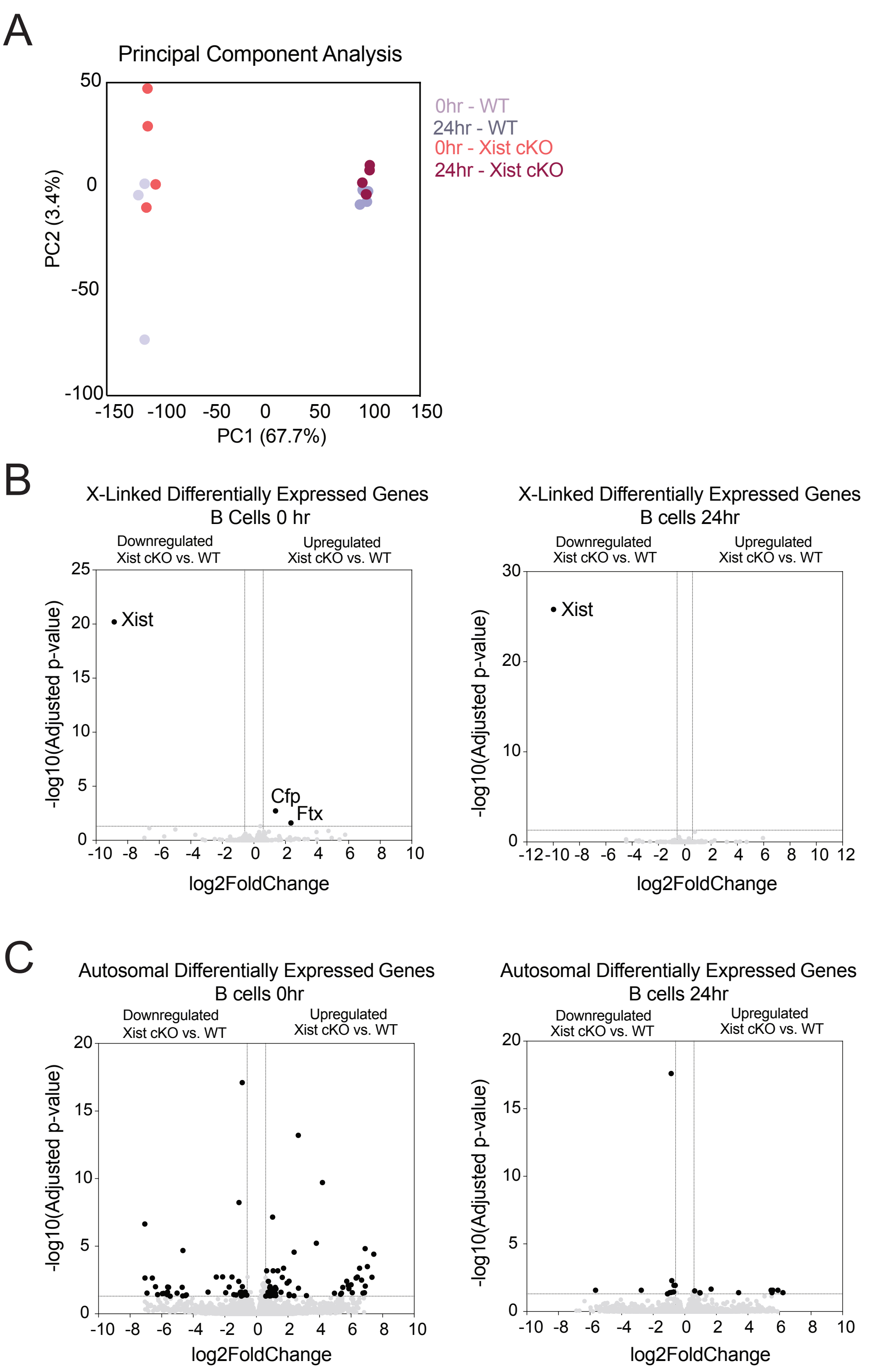

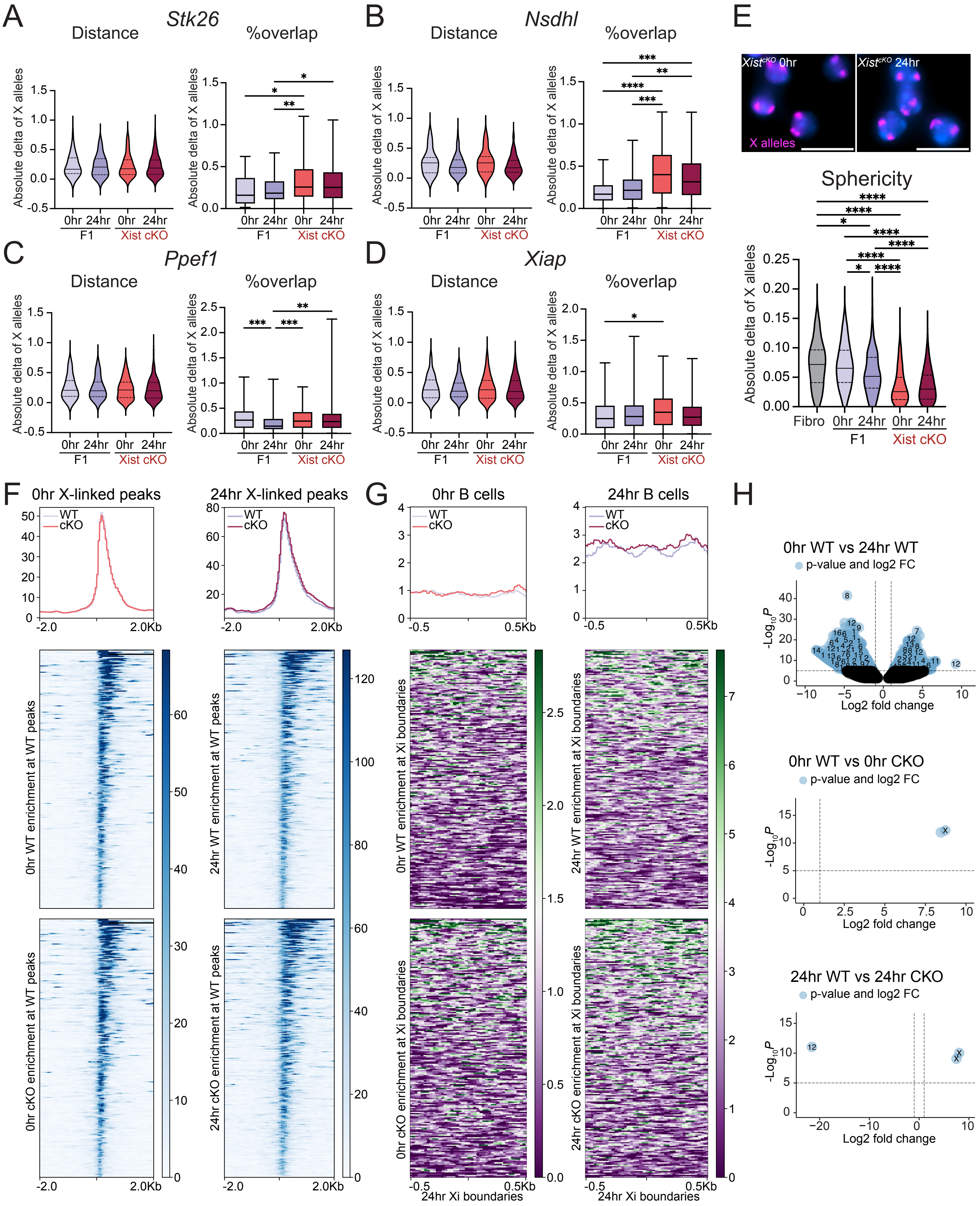
